# Histone modifications define distinct nucleosome positions in cell-free DNA

**DOI:** 10.64898/2026.10.26.761423

**Authors:** Yukun Ou, Gunsagar S. Gulati, Caroline F. Tyndall, Medha Pandey, Ji-Heui Seo, Ze Zhang, Zhenjie Jin, Garyoung Gary Lee, Rashad Nawfal, Taylor A. Hoggood, Sadia D. Shahbazi, John Ludwig, Ziwei Zhang, Elio Ibrahim, Gaelle Nafeh, Jacob E. Berchuck, Adam G. Sowalsky, Fatima Karzai, Matthew L. Freedman, David Y. Takeda, Sylvan C. Baca

## Abstract

Circulating cell-free DNA (cfDNA) fragmentation patterns reflect nucleosome positioning. We asked whether variation in nucleosome positions in cfDNA can reveal chromatin states and associated histone modifications. Using cell-free chromatin immunoprecipitation sequencing (cfChIP-seq) in plasma from patients with prostate cancer, we identified 35,120 modification-associated positions (MAPs), loci where nucleosomes bearing active (H3K4me2) and repressive (H3K9me3) histone marks occupy reproducibly offset positions. These offsets recurred across histone marks, an independent cfChIP-seq cohort, and nuclease-digested chromatin from cell lines. MAPs were enriched in transcribed genes but depleted near transcription start sites and CTCF-binding sites where nucleosome positioning is constrained. Across 2,535 cfDNA sequencing datasets, nucleosome protection at a subset of MAPs correlated with tumor fraction (TF), reflecting differences in histone modification status between cancer-derived and non-malignant chromatin. In targeted sequencing panels, a 50-MAP model using fragments from < 0.001% of the genome distinguished samples with tumor fractions ≥3% vs 0% (AUCs = 0.91-0.96 across prostate, breast, and lung cancers) and outperformed a state-of-the-art fragmentomics-based tumor fraction predictor. MAP-derived estimates correlated with reference TFs (*r* = 0.89) and longitudinal changes in TF (*r* = 0.91). MAP protection at ASCL1 binding sites identified neuroendocrine transformation of prostate cancer from cfDNA. These findings establish histone-modification-associated nucleosome positioning as a mutation-independent readout of tumor fraction and cancer state from routine cfDNA sequencing.

## INTRODUCTION

Circulating cell-free DNA (cfDNA) carries information about the chromatin of the cells from which it originates. Histones and other DNA-bound proteins protect DNA from cleavage, shaping cfDNA fragment lengths, end motifs, and genomic coverage and leaving footprints of tissue-specific chromatin organization in circulating DNA.^1–3^ Nucleosome organization around transcription start sites, enhancers, and transcription-factor binding sites has consequently been used to infer genomic activity, distinguish cancer types, and predict chronological age.^4–12^ However, conventional cfDNA sequencing does not directly reveal the chromatin states associated with individual nucleosome footprints.

Histone modifications provide a molecular record of chromatin state.^13–17^ For example, H3K4me2 and H3K9me3 are enriched in active and repressive chromatin, respectively.^17,18^ Cell-free chromatin immunoprecipitation with sequencing (cfChIP-seq) can resolve these states noninvasively in patients by enriching circulating nucleosomes marked by specific histone modifications, revealing epigenomic programs and their dynamic regulation in health and disease.^19–22^ However, each modification requires a separate antibody-based enrichment and therefore cannot be directly resolved from conventional whole-genome or targeted sequencing of total cfDNA. Histone modifications have also been associated with differences in cfDNA fragment lengths and end motifs,^10,23–25^ providing indirect readouts of chromatin state and raising the possibility that other features of nucleosome organization may encode similar information.

Nucleosomes can occupy highly reproducible positions that can be mapped to within a few base pairs and are shaped by DNA sequence, chromatin remodelers, and bound factors.^26–37^ These positions can shift with transcriptional activity, differentiation, aging, and malignant transformation.^12,38–41^ We hypothesized that circulating nucleosomes carrying different histone modifications preferentially occupy distinct positions within a genomic locus. If preserved in circulating DNA, then measuring these positional differences could identify cancer signals, without performing immunoprecipitation. Pairing an active and a repressive mark at the same locus serves two purposes. Where the two marks occupy different positions, the position a cfDNA fragment protects indicates the chromatin state of the nucleosome it came from. Because both positions lie within one nucleosome footprint, the difference in protection between them is measured within a single locus, so local coverage and copy number affect both positions alike. In plasma, which mixes DNA from tumor and blood cells, a locus at which the two cell types carry different marks would therefore report the tumor contribution.

Here, we identify modification-associated positions (MAPs), genomic loci at which circulating nucleosomes marked by active (H3K4me2) and repressive (H3K9me3) histone modifications occupy reproducibly distinct positions (**Figure 1A**). We show that MAPs validate in an independent plasma cohort, extend to additional histone modifications, and are detectable in non-immunoprecipitated cfDNA and cellular chromatin. We further show that differential nucleosome protection at MAPs provides a compact readout of tumor fraction from whole-genome and targeted cfDNA sequencing (**Figure 1B**). MAPs can be measured at predefined loci without genome-wide sequencing,^42,43^ and do not depend on tumor mutation burden or copy-number status,^42,44^ which vary across cancer types. Using small MAP panels, we estimate tumor fraction across independent cohorts, multiple cancer types, and serial plasma samples, establishing histone-associated nucleosome positioning as a mutation-independent source of information for cancer profiling from cfDNA.

**Figure 1.**
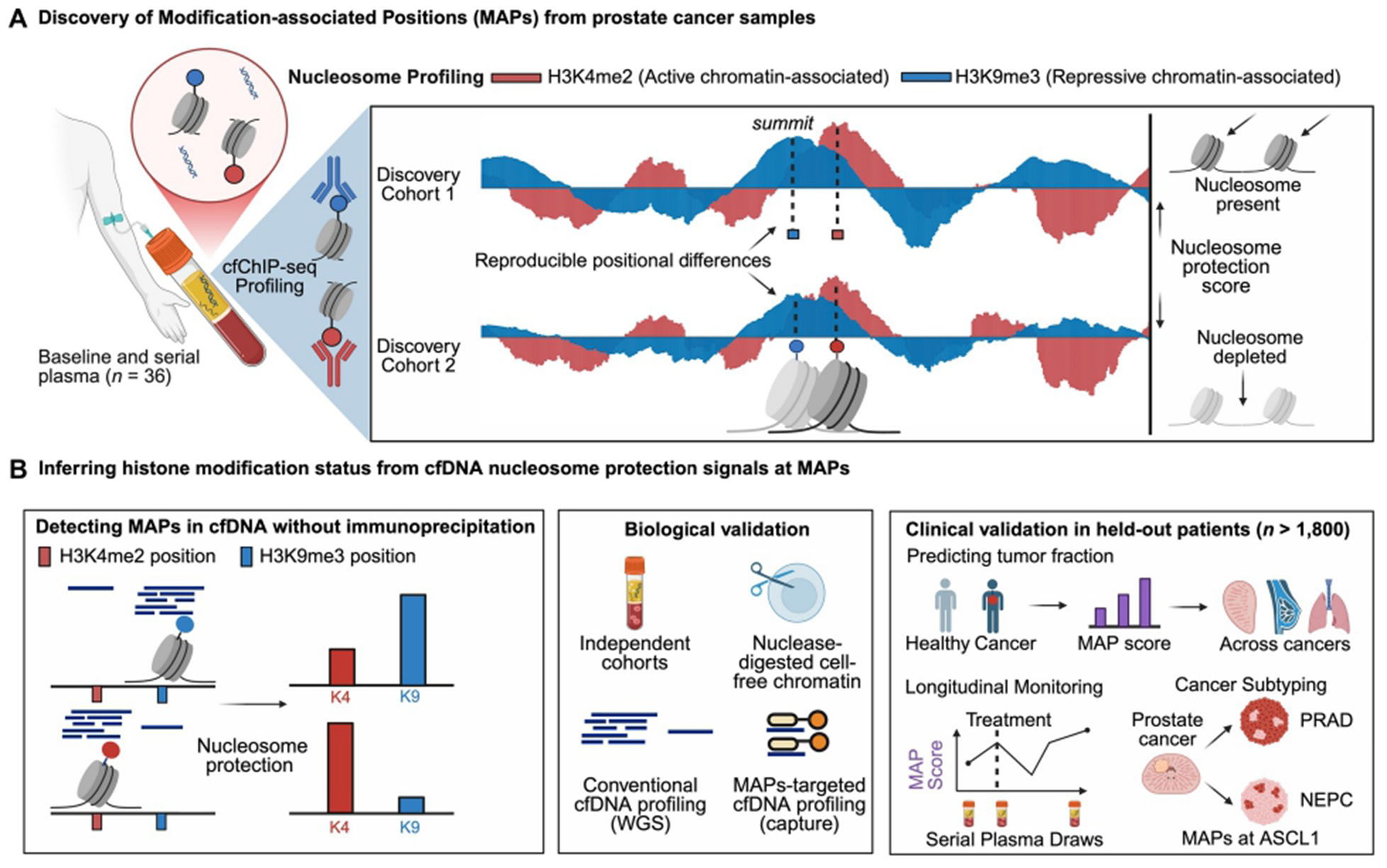
Overview of histone modification-associated positions (MAPs). **(A)** Discovery of MAPs from H3K4me2 (red; active chromatin) and H3K9me3 (blue; repressive chromatin) cfChIP-seq of baseline and serial plasma from patients with prostate cancer in two discovery cohorts (36 specimens; cohort 1, 7 patients and 17 specimens; cohort 2, 8 patients and 19 specimens). The probabilistic nucleosome score (PNS) is positive where a nucleosome is present and negative where nucleosomes are depleted. Dashed lines mark the PNS summit of each mark, the inferred nucleosome center (dyad; squares); the two marks peak at reproducibly different positions in both cohorts. **(B)** Inferring histone modification status at MAPs from cfDNA. Left, in cfDNA sequenced without immunoprecipitation, the relative protection of fragments at the H3K4me2- and H3K9me3-associated positions of a MAP indicates which nucleosome position, and hence which chromatin state, predominates. Middle, biological validation in an independent cfChIP-seq cohort, in nuclease-digested chromatin from prostate cancer cells, and in unenriched cfDNA sequenced genome-wide (WGS) or by hybrid capture of MAPs. Right, clinical validation (prostate cancer, 1,197 patients; breast cancer, 336; lung cancer, 322; **Methods**): tumor fraction prediction across cancer types, longitudinal monitoring of serial plasma draws, and cancer subtyping of prostate adenocarcinoma (PRAD) and neuroendocrine prostate cancer (NEPC) from MAPs at ASCL1 binding sites.

## RESULTS

### Paired H3K4me2 and H3K9me3 positions define MAPs

We first asked whether nucleosomes bearing active and repressive histone modifications occupy distinct positions at the same genomic locus in plasma. In two discovery cohorts of patients with metastatic prostate cancer followed by serial plasma cfChIP-seq (cohort 1, 7 patients and 17 specimens; cohort 2, 8 patients and 19 specimens; 36 specimens profiled for both marks; **Figure 1A**; **Supplementary Table 1**), we inferred nucleosome centers (dyads) independently from H3K4me2- and H3K9me3-associated fragments, pooling the fragments of each mark across the specimens of each cohort. We retained high-confidence dyads identified by two complementary nucleosome-positioning methods, the probabilistic nucleosome score (PNS) and DANPOS3 (**Figure S1A–C**; **Figure S2A**).^45,46^ Within each cohort, H3K4me2- and H3K9me3-associated dyads were paired when they were reciprocal nearest neighbors within 73 bp, approximately half the 147 bp of DNA wrapped around a nucleosome core.^47,48^ Dyads separated by this distance therefore share more than half of their underlying nucleosomal DNA, consistent with alternative positions of overlapping nucleosomes rather than distinct, adjacent nucleosomes (**Figure S1F**; **Supplementary Note 1**).

We defined modification-associated positions (MAPs) as loci at which H3K4me2- and H3K9me3-associated dyads were reproducibly separated by 10–73 bp across both cohorts and maintained the same upstream mark. As a comparison class, copositioned K4/K9 anchors were separate loci at which the two dyads were separated by only 0–4 bp in both cohorts (**Figure S1E,F**; **Methods**). The 10-bp lower bound corresponds approximately to one helical turn of nucleosomal DNA, consistent with alternative positions that preserve rotational setting, whereas the copositioned cutoff falls within half a turn.^29,47^ Among 120,489 loci with consistent dyad positions across cohorts, we identified 35,120 MAPs and 4,774 copositioned anchors (**Figure S1A**). The median separation at MAPs, averaged over the two discovery cohorts, was 28 bp (interquartile range, 20.5–38 bp).

MAPs exhibited distinct genomic properties compared with copositioned anchors, including reduced enrichment within 1 kb of transcription start sites (TSSs; **Figure S1G–I**) and at CTCF-binding sites identified in the prostate cancer cell line LNCaP (**Supplementary Table 2**). These differences persisted after stratifying loci on total cfChIP-seq fragments and varying the distance threshold used to define copositioned anchors (2–8 bp). This pattern is consistent with prior evidence that transcriptional regulatory regions and CTCF-binding sites exhibit strongly organized nucleosome positioning,^26,35^ suggesting that modification-associated positional differences may preferentially arise at genomic loci with fewer constraints on nucleosome positioning. Supporting the robustness of these measurements to sampling bias, PNS peaks were recovered in 89–94% of eligible profiles after randomly removing one-quarter of fragments (**Figure S1J**). These analyses establish a reproducible set of MAPs representing distinct H3K4me2- and H3K9me3-associated nucleosome positions in plasma.

### Independent cfChIP-seq profiles recover MAP separation and orientation

To determine whether MAPs represent reproducible nucleosome positions, we evaluated their separation and orientation in independent cfChIP-seq datasets. In a published cohort of ten plasma specimens from patients with advanced cancer (independent cohort; Sipola et al.^21^), H3K4me2- and H3K9me3-associated positions showed progressively greater separation across copositioned anchors and the two MAP groups defined by increasing discovery displacement (**Figure 2A–D**). Median separation in the independent cohort was 1 bp at copositioned anchors and 19 bp and 54.5 bp in the two MAP groups, closely recapitulating the original spacing defined in the discovery cohorts (25 and 47 bp; **Figure 2B**). Moreover, the relative orientation of H3K4me2- and H3K9me3-marked nucleosomes was preserved at 71% and 75% of measurable MAPs, substantially exceeding that expected by chance (**Figure 2E**).

**Figure 2.**
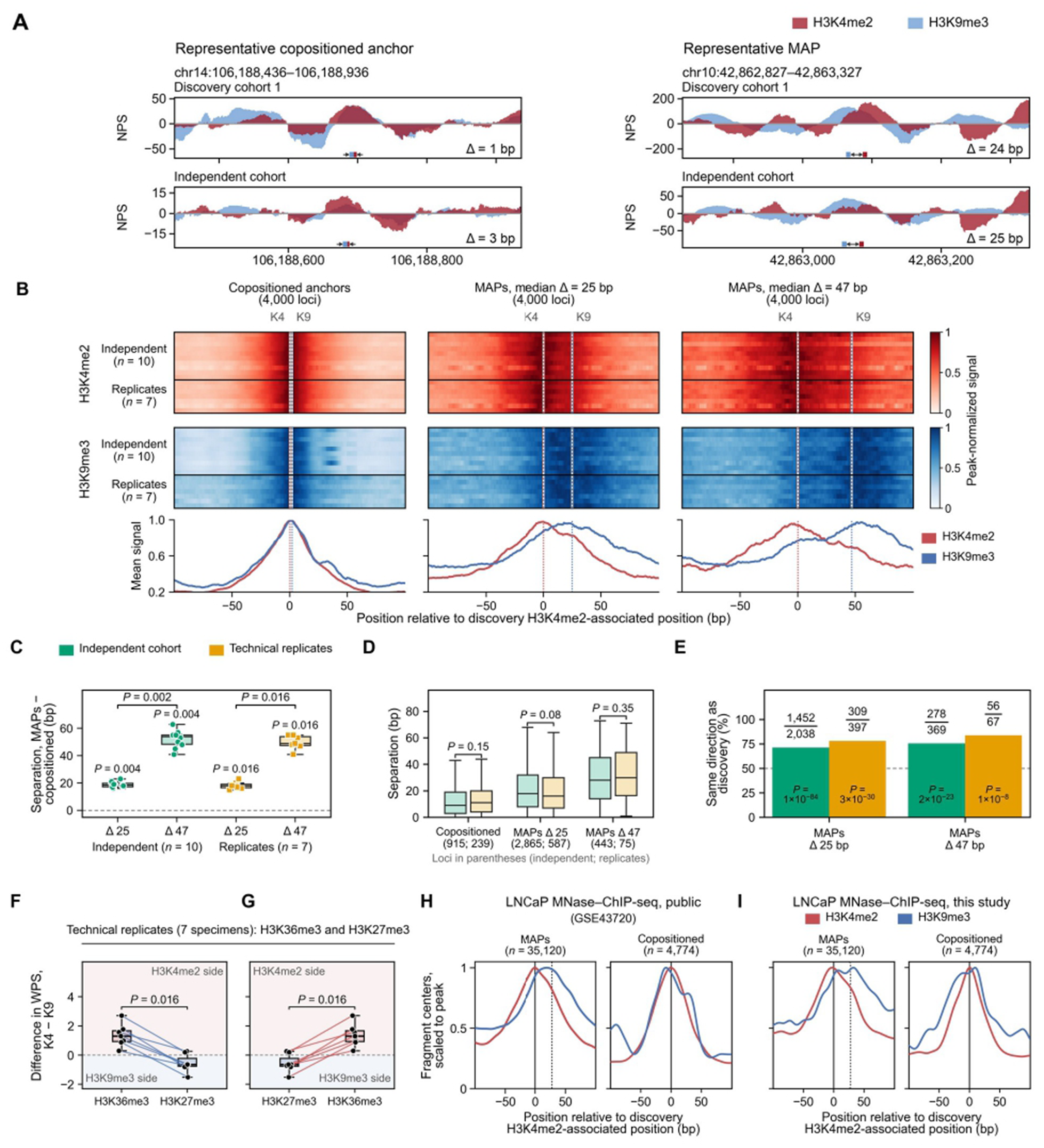
MAPs show consistent offsets of active vs. inactive nucleosomes across cohorts and histone marks. **(A)** Representative copositioned anchor (left) and MAP (right): probabilistic nucleosome score (PNS; unitless) of H3K4me2 (red) and H3K9me3 (blue) cfChIP-seq in discovery cohort 1 and in the independent cohort (Sipola et al.^21^). Overlapping signal appears as a mixed color. Squares mark the inferred dyads (PNS maxima), and Δ is the distance between the H3K4me2 and H3K9me3 dyads. **(B)** H3K4me2 (top) and H3K9me3 (bottom) profiles at 4,000 loci drawn at random from each of three groups: copositioned anchors and the MAPs 10–40 bp and more than 40 bp apart (Δ, median separation averaged over the two discovery cohorts; **Methods**).

To account for technical bias, the H3K4me2 and H3K9me3 cfChIP-seq libraries were independently generated by the Takeda and Wyatt laboratories from seven of these plasma specimens (inter-laboratory replicates). These replicates from different institutions reproduced both the separation and orientation of MAPs, with directional concordance of 78% and 84% across the two MAP groups (**Figure 2B–E**), and separation at individual loci did not differ significantly between the independent cohort and its inter-laboratory replicates (Figure 2D). Similar patterns were observed across 71,504 discovery-paired loci in pooled validation profiles (**Figure S3**). These results demonstrate that MAP spacing and orientation are reproducible across independent plasma specimens, cfChIP-seq datasets, and laboratories.

### MAP positions distinguish active and repressive chromatin states

We next tested whether MAP positions generalized to histone modifications beyond the H3K4me2 and H3K9me3 marks used for their discovery. Using H3K36me3, a mark associated with transcription, and H3K27me3, a mark of Polycomb-repressed chromatin,^49^ we compared, at each MAP, the protection of the H3K4me2-associated position with that of the H3K9me3-associated position, measured by the windowed protection score (WPS), and subtracted the same comparison in the unenriched input library from the same plasma (Methods). After this correction, H3K36me3 preferentially protected the H3K4me2-associated position in all seven specimens, whereas H3K27me3 preferentially protected the H3K9me3-associated position in five (**Figure 2F,G**; **Figure S4**). Across all seven specimens, the relative positional preference of H3K36me3 was shifted toward the H3K4me2-associated position compared with H3K27me3 (two-sided sign test, *P* = 0.016). Thus, the two positions defined by MAPs distinguish nucleosomes associated with broader states of active and repressive chromatin rather than being specific to the histone marks used for their discovery.

### Prostate cancer cell MNase–ChIP-seq recapitulates plasma-defined MAP positions

To determine whether plasma-defined MAPs reflect nucleosome positions present in prostate cancer cells, rather than features introduced by plasma fragmentation or immunoprecipitation, we examined H3K4me2 and H3K9me3 micrococcal nuclease (MNase)–ChIP-seq, in which nuclease digestion preserves nucleosome-protected DNA and provides an orthogonal readout of nucleosome positioning. In published LNCaP prostate cancer profiles,^50^ H3K4me2- and H3K9me3-associated fragments separated toward their respective plasma-defined MAP positions, whereas both marks remained centered at copositioned anchors (**Figure 2H**). We then generated H3K4me2 and H3K9me3 MNase–ChIP-seq profiles from LNCaP cells and reproduced the same modification-associated separation, while copositioned anchors remained aligned (**Figure 2I**). Thus, nucleosome-protected chromatin from prostate cancer cells recapitulates the modification-associated positions identified in prostate cancer plasma, supporting MAPs as physical differences in nucleosome positioning.

Each row is one sample: ten from the independent cohort and seven inter-laboratory replicates, new libraries prepared from seven of the same ten plasma specimens. Profiles are aligned to the discovery H3K4me2-associated position and scaled to their peak; dashed lines mark the discovery H3K4me2- and H3K9me3-associated positions. Lower plots show the mean of the 17 rows. **(C)** Separation in each MAP group minus that at copositioned anchors, one point per specimen. *P values above each group, one-sided exact sign test that separation at MAPs exceeds that at copositioned anchors, Holm-corrected across the four comparisons;* brackets, the two MAP groups compared within the independent cohort and within the inter-laboratory replicates (two-sided exact sign test). **(D)** Separation between the H3K4me2 and H3K9me3 dyads at individual loci; brackets compare the independent cohort with its inter-laboratory replicates (two-sided Mann–Whitney test). **(E)** Percentage of measurable MAPs at which the two positions keep the discovery orientation; numbers above the bars give loci with the same orientation over measurable loci; *P, one-sided exact binomial test against 50%; dashed line, 50%. (F,G)* Difference in windowed protection score (WPS) between the H3K4me2- and H3K9me3-associated positions for H3K36me3 and H3K27me3 cfChIP-seq of the seven inter-laboratory replicate specimens, after subtracting the same difference in the input library of each specimen and expressed per 100 fragments, shown from the H3K36me3 (F) and H3K27me3 (G) sides. Lines connect the same specimen; *P*, two-sided exact sign test. **(H)** Public LNCaP MNase–ChIP-seq (Chen et al.^50^; GSE43720; two replicates per mark, averaged). Lines show the density of fragment centers around the 35,120 MAPs (left) and the 4,774 copositioned anchors (right), oriented toward the H3K9me3-associated position, smoothed and scaled to their peak; solid line, discovery H3K4me2-associated position; dotted line, median discovery H3K9me3-associated position of MAPs (+28 bp). Fragment centers were placed 73 bp from the 5′ end of each 35-bp single-end read. **(I)** MNase–ChIP-seq of LNCaP cells generated in this study (centers of 120–180-bp paired-end fragments; representative of three independent experiments), displayed as in H. Box plots show the median, interquartile range (IQR) and whiskers to the most extreme values within 1.5 × IQR.

### MAPs preferentially occur in regions with less constrained nucleosome positioning

To identify genomic features associated with MAPs, we compared MAPs with copositioned anchors, focusing on features that might influence nucleosome positioning. MAPs were broadly distributed across the genome (**Figure S2B**) and had similar mappability and GC content to copositioned anchors (**Figure S2C,D**). They were less frequent near transcription start sites and promoter chromatin states and more frequent within gene bodies, transcribed regions, quiescent chromatin, and distal enhancer-like regions (**Figure 3A**; **Figure S5A–D**). MAPs were also more likely to occur within genes expressed across tissues or in prostate (**Figure 3B**). Because copositioned anchors had more cfChIP-seq fragments than MAPs, subsequent comparisons were stratified on H3K4me2 and H3K9me3 read depth (**Methods**).

**Figure 3.**
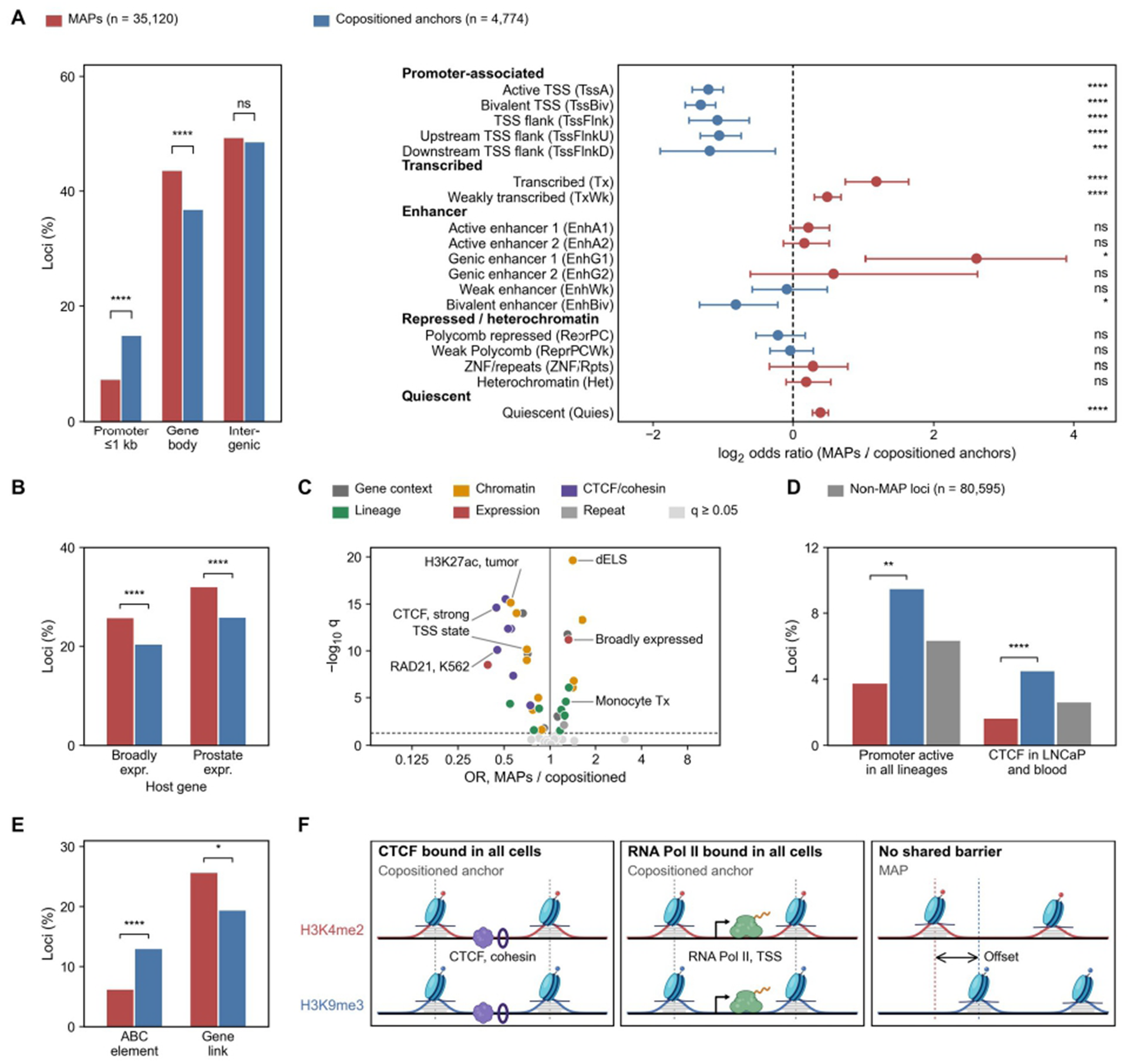
MAPs preferentially occur in regions with less constrained nucleosome positioning. **(A)** Left, percentage of 35,120 MAPs (red) and 4,774 copositioned anchors (blue) within 1 kb of a transcription start site (TSS) and overlapping gene bodies or intergenic regions. Right, log2 odds ratios (MAPs versus copositioned anchors) for the 18 EpiMap chromatin states of normal prostate gland tissue (BSS01456), with chromosome-bootstrap 95% confidence intervals; stars, two-sided Fisher’s exact test *P* < 0.05, 0.01, 0.001 and 0.0001. **(B)** Percentage of loci in the body of a host gene (the protein-coding gene whose span contains the midpoint, more than 1 kb from any coding TSS) expressed (GTEx median TPM ≥1) in at least 40 of 54 tissues or in prostate. **(C)** Mantel–Haenszel odds ratio (MAPs versus copositioned anchors) and Benjamini–Hochberg q value for 59 reference region sets including gene context (distance to the nearest TSS, CpG islands, gene bodies and intergenic regions), chromatin states, SCREEN candidate cis-regulatory elements, H3K27ac in prostate tumors and normal prostate, LNCaP ABC elements, CAGE, CTCF and cohesin peaks, prostate lineage factors, lineage activity, host-gene expression and repeats (**Supplementary Table 2**). Colors, set families for sets with q < 0.05; light gray, q ≥ 0.05; dashed line, q = 0.05. dELS, distal enhancer-like signature; Tx, transcribed state. **(D)** Percentage of loci at a promoter in a TSS-associated state in prostate, neutrophils and monocytes, and near a CTCF peak bound in LNCaP and in K562, GM12878 or neutrophils, for MAPs, copositioned anchors and Non-MAP loci (gray, shown for reference); P, Wald test from one logistic regression of MAPs versus copositioned anchors on both features and on chromatin activity restricted to one or two of the three cell types. In B and C, comparisons are stratified on H3K4me2 and H3K9me3 read depth (16 strata; Cochran–Mantel–Haenszel test); stars, q or *P* < 0.05, 0.01, 0.001 and 0.0001. **(E)** Percentage of loci at LNCaP Activity-by-Contact (ABC) candidate elements defined by the ABC model and percentage with a predicted enhancer–gene link among loci at distal candidate elements; *P*, two-sided Fisher’s exact test. **(F)** Proposed model for MAPs. Where factors such as CTCF (left) or RNA polymerase II (middle) are bound in all contributing cells, nucleosomes carrying either mark share positions (copositioned anchors). Without such a shared barrier, H3K4me2- and H3K9me3-marked nucleosomes may occupy offset positions (MAPs, right).

The strongest differences involved factors associated with constrained nucleosome organization. CTCF and cohesin binding sites are surrounded by regularly positioned nucleosome arrays.^35,51,52^ Across reference genomic annotations, CTCF and cohesin sites were among the most strongly depleted features near MAPs relative to copositioned anchors (**Figure 3C**; **Supplementary Table 2**). This depletion was most apparent at CTCF sites and promoters shared across prostate and blood lineages, whereas lineage-restricted chromatin activity did not distinguish MAPs from copositioned anchors (**Figure 3D**). Moreover, CTCF and cohesin binding became progressively less frequent as the separation between H3K4me2- and H3K9me3-associated positions increased, even after adjustment for genomic features and coverage (**Supplementary Table 3**; **Figure S5B**). These findings suggest that MAPs preferentially arise at loci where nucleosome positions are less constrained by stable chromatin-organizing features.

Given the enrichment of MAPs in distal and enhancer-like chromatin, we next asked whether these loci were associated with regulatory function. MAPs were less frequent than copositioned anchors at H3K27ac-marked regions and at LNCaP candidate regulatory elements identified by the Activity-by-Contact (ABC) model,^53^ indicating that MAPs are not simply a feature of active enhancers (**Figure 3E**; **Figure S5E**; **Supplementary Table 4**). Together with the depletion at shared CTCF sites and promoters, these comparisons support a model in which barriers present in every contributing cell, such as bound CTCF and cohesin or promoter-bound RNA polymerase II, fix nucleosome positions regardless of chromatin state, whereas away from such barriers nucleosomes in active and repressive chromatin can occupy different positions (**Figure 3F**).

### Nucleosome positions at a subset of MAPs associate with tumor-versus blood-derived cfDNA

We asked whether the two positions at a MAP could reflect nucleosomes originating from different cell populations, given that plasma from patients with cancer contains both blood- and tumor-derived cfDNA.^1,54^ If so, the relative cfChIP-seq signals at the two positions should change in opposite directions with increasing tumor fraction. Across the 36 discovery specimens, 3,395 of the 35,120 MAPs showed this pattern. At 1,776 MAPs, H3K4me2 signal increased and H3K9me3 signal decreased with tumor fraction (K4/TF-correlated MAPs), whereas 1,619 showed the reverse pattern (K9/TF-correlated MAPs; **Figure 4A**; **Supplementary Table 5**).

**Figure 4.**
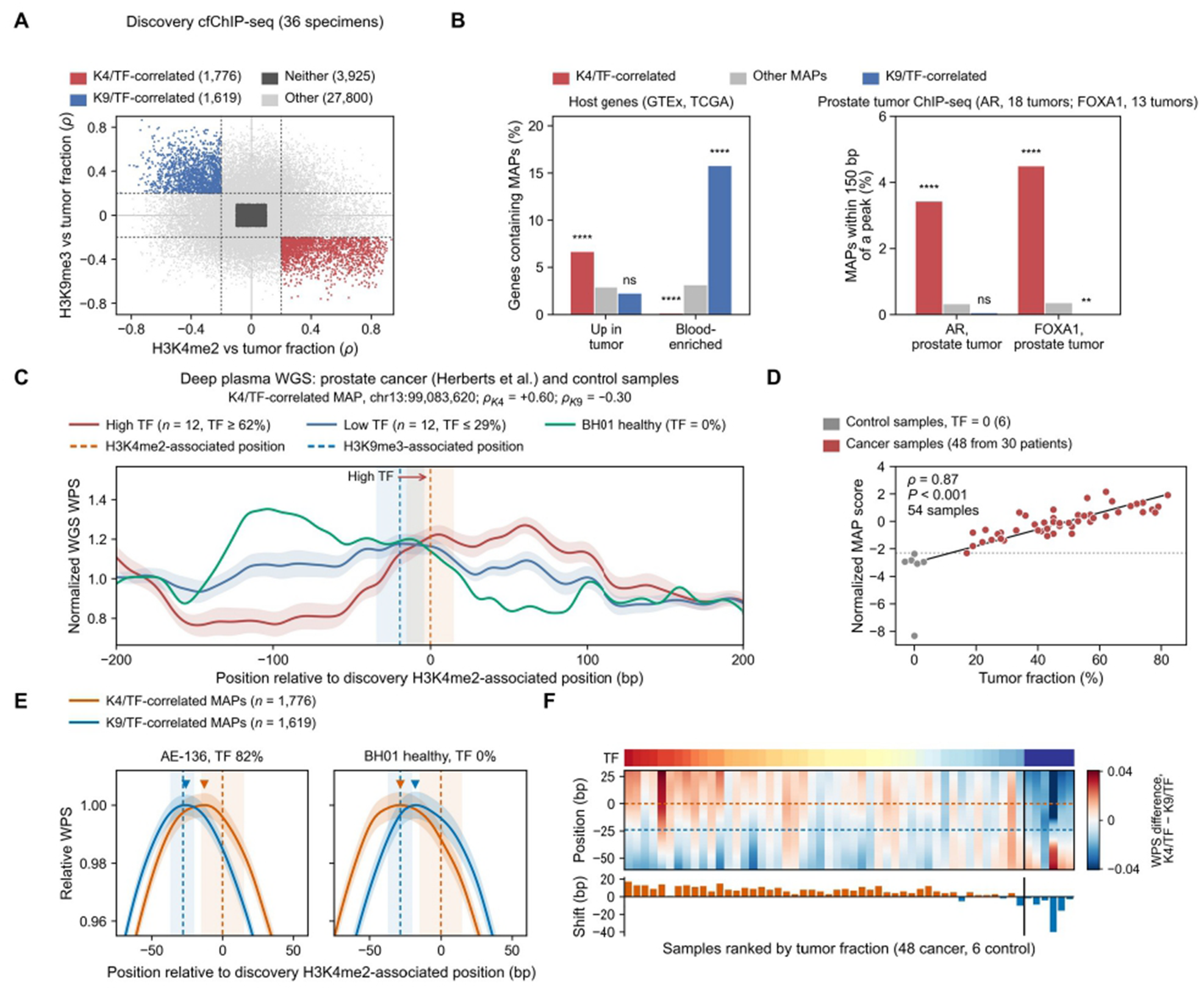
Protection at fixed MAP positions follows tumor fraction in whole-genome cfDNA sequencing. **(A)** Selection of tumor fraction-correlated MAPs. Spearman correlation of H3K4me2 (x) and H3K9me3 (y) cfChIP-seq signal with tumor fraction (TF) across 36 discovery specimens at all 35,120 MAPs; dashed lines, |ρ| = 0.2, a cutoff at which opposite-direction coupling is more frequent at MAPs than at matched Non-MAP loci (**Methods**; **Figure S6F**). Red, K4/TF-correlated MAPs (*n* = 1,776); blue, K9/TF-correlated MAPs (*n* = 1,619); dark gray, MAPs at which neither signal correlated with TF (|ρ| < 0.1 for both marks; *n* = 3,925); light gray, other MAPs. **(B)** Left, percentage of genes containing MAPs (host genes: protein-coding genes whose transcribed region contains the MAP) that are upregulated in primary prostate tumors (TCGA) or blood-enriched (GTEx). Right, percentage of MAPs within 150 bp, about one nucleosome, of an AR or FOXA1 peak in prostate tumors. Red, K4/TF-correlated; gray, other; blue, K9/TF-correlated MAPs; stars, two-sided Fisher’s exact test against other MAPs. **(C)** WGS protection at a K4/TF-correlated MAP in the quartiles of the 48 cancer specimens with the highest TF (12 specimens, TF ≥62%) and the lowest TF (12 specimens, TF ≤29%), mean ± s.e.m., and in BH01; dashed lines, the H3K4me2-associated (orange) and H3K9me3-associated (blue) positions from discovery cfChIP-seq, ±15 bp shaded; ρK4 and ρK9, Spearman correlations of H3K4me2 and H3K9me3 cfChIP-seq signal with TF in discovery. **(D)** Normalized MAP score over the 8,044 MAPs from 48 prostate cancer cfDNA datasets from Herberts et al.^44^ (red) and six control samples (gray; offset horizontally). Dashed line, the lowest cancer plasma MAP score; solid line, least-squares fit; ρ, Spearman correlation across the 54 samples; *P*, two-sided permutation test exchanging patients as whole blocks, with each control sample as its own block (10,000 permutations; **Methods**). **(E)** Composite WPS over K4/TF-correlated (orange) and K9/TF-correlated (blue) MAPs, each scaled to its peak, in a specimen with 82% TF and in a healthy control sample; shading around each composite, ±1 s.e.m. across MAPs; triangles, relative WPS peaks; dashed lines, the H3K4me2-associated position and the median H3K9me3-associated position (−28 bp), with ±15 bp around the H3K4me2-associated position and the interquartile range of the H3K9me3-associated position (−37 to −20 bp) shaded. **(F)** For every sample (columns, ranked by TF), the K4/TF-correlated composite minus the K9/TF-correlated composite is shown (color scale, WPS difference). Dashed lines show the H3K4me2-associated position (orange) and the median H3K9me3-associated position (blue); lower track, peak shift (K4/TF-correlated minus K9/TF-correlated peak position, bp). WPS, windowed protection score.

The genomic contexts of these MAPs supported distinct cellular origins. Genes containing K4/TF-correlated MAPs were enriched for genes upregulated in primary prostate tumors, and these MAPs were more frequently located near AR and FOXA1 binding sites in prostate tumors. In contrast, genes containing K9/TF-correlated MAPs were strongly enriched for blood-enriched genes (expressed in whole blood at least fourfold above the mean of other tissues in GTEx) (**Figure 4B**). These associations suggest that, at a subset of MAPs, the two histone-mark-associated nucleosome positions capture differences between tumor- and blood-derived chromatin.

### Nucleosome protection at MAPs tracks tumor fraction in whole-genome cfDNA sequencing

We next asked whether the tumor-associated changes at MAPs could be detected directly from cfDNA sequenced without immunoprecipitation. We analyzed published deep whole-genome sequencing (WGS) of 48 cfDNA samples from 30 patients with metastatic castration-resistant prostate cancer (mCRPC; tumor fraction, 17–82%; Herberts cohort)^44^ and six control samples without detectable tumor DNA. We calculated fragment-protection profiles around the fixed MAP positions and compared nucleosome protection at the H3K4me2- and H3K9me3-associated positions (**Figure 4C**; **Methods**). At a representative K4/TF-correlated MAP, specimens with high tumor fractions showed greater protection at the H3K4me2-associated position and less at the H3K9me3-associated position than specimens with low tumor fractions and healthy control cfDNA (**Figure 4C**). We therefore asked whether this tumor fraction-associated pattern was reproducible across MAPs. We aggregated protection across 8,044 MAPs selected and oriented using only the discovery cfChIP-seq data, such that increasing scores reflected the expected tumor-associated shift in protection. The resulting MAP score separated all six controls from the 48 cancer specimens and increased strongly with tumor fraction (Spearman ρ = 0.87; *P* < 0.001; **Figure 4D**). The association remained strong among cancer specimens alone (ρ = 0.82), across patient means, and within patients with serial samples, and persisted after adjustment for sequencing depth and fragment size (**Figure S6A–D**). Aggregation substantially strengthened the relationship with tumor fraction relative to individual MAPs, whose correlations were generally weak (**Figure S6E**).

Tumor fraction was also associated with a shift in the position of maximal nucleosome protection. Across K4/TF-correlated MAPs, protection shifted toward the H3K4me2-associated position in 45 of 48 cancer specimens but in none of the six control profiles, and the difference between K4/TF- and K9/TF-correlated MAPs increased with tumor fraction (**Figure 4E,F**). This relationship was robust to the correlation threshold used to define tumor-associated MAPs (**Figure S6F**). Thus, increasing tumor fraction altered not only the relative protection at the two MAP positions but also the inferred position of the protected nucleosome.

Because fragment-protection measurements depend on the number of observed fragments, performance decreased at lower sequencing depth. Discrimination between specimens with low and high tumor fractions declined substantially with downsampling (**Figure S7A–E**), while in silico mixtures into deeply sequenced healthy cfDNA raised the MAP score steadily from 0 to 20% tumor fraction (**Figure S7F**). These results show that histone-mark-associated nucleosome positions identified by cfChIP-seq leave measurable protection signatures in conventional cfDNA sequencing that reflect both tumor fraction and its longitudinal changes.

### MAP signals in existing targeted sequencing data

We next asked whether MAP signals could be discerned in other cohorts that used targeted sequencing approaches. We first examined the Emory capture cohort, comprising hybrid-capture sequencing of plasma from 16 patients with mCRPC and seven cancer-free controls. The capture panel was designed for a separate project, and 224 of the 8,044 MAPs defined from the discovery cfChIP-seq data lay within its targets. At these MAPs with at least ten fragments at both positions, we measured the difference in fragment protection between the two MAP positions after normalization to local coverage and combined these differences into a MAP score. Scores were higher in patients with mCRPC than in controls (AUC = 1.00; P = 8 × 10^−6^; Figure 5A). Representative loci showed the expected differences in protection between mCRPC and control plasma (**Figure 5B**; **Figure S8**). Thus, tumor fraction-associated signals at MAPs can be measured from cfDNA enriched by hybrid capture.

**Figure 5.**
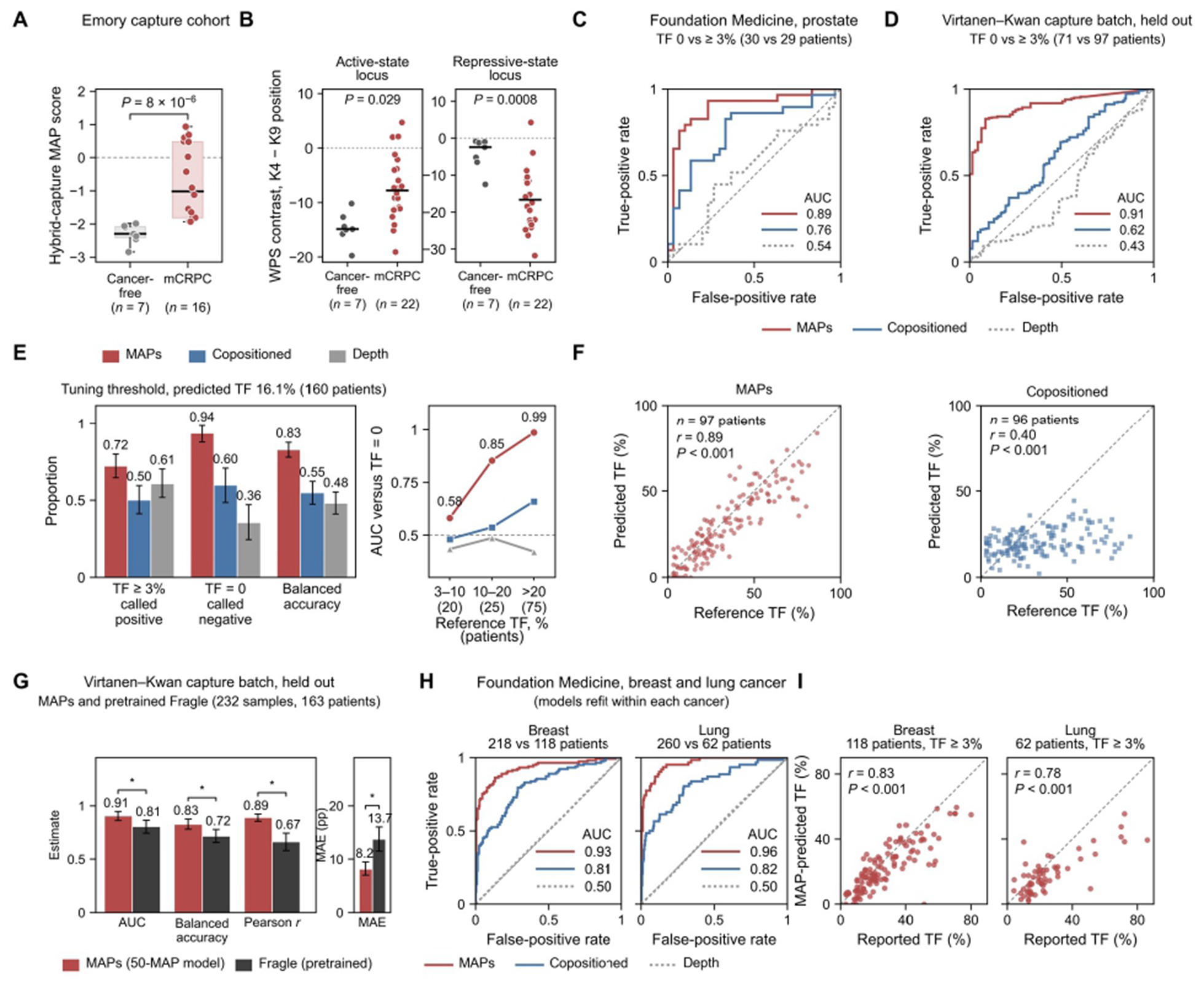
MAPs on targeted sequencing panels estimate tumor fraction across cohorts and cancer types. **(A)** Emory capture cohort: hybrid-capture MAP score at the 224 MAPs of Figure 4D that lie within the capture targets, each with at least ten fragments at both positions for cancer-free controls (*n* = 7) and patients with metastatic castration-resistant prostate cancer (mCRPC; *n* = 16); one point per patient; two-sided Mann–Whitney test. **(B)** Windowed protection score (WPS) contrast between the H3K4me2- and H3K9me3-associated positions at two representative loci at which protection is expected to shift toward the H3K4me2-associated position (active-state locus) or the H3K9me3-associated position (repressive-state locus) with tumor DNA; one point per specimen (7 cancer-free, 22 mCRPC); black lines, medians; *P*, patient-label permutation test (13 patients with mCRPC and seven controls). **(C)** Foundation Medicine cohort, prostate cancer: ROC curves for reported tumor fraction (TF) 0 versus ≥3% in 59 patients (30 and 29) from nested cross-validation with identical folds for models built on 1,342 MAPs (red), 412 copositioned anchors (blue) and sequencing depth (gray dotted); the model on 16 fragment-length and sequencing features is compared in **Figure S9I**. **(D)** Virtanen–Kwan cohort,^55^ capture batch (held-out set of 232 samples from 163 patients): ROC curves for reference TF 0 versus ≥3% for the 50-MAP model and for models built on copositioned anchors or sequencing depth, each fitted on development patients and applied unchanged. Reference TF is the source study’s ctDNA fraction. **(E)** The 50-MAP model of D at the threshold that maximized balanced accuracy in tuning patients: share of patients with TF ≥3% called positive, share with TF 0 called negative, and balanced accuracy, with 95% patient-bootstrap confidence intervals; right, AUC for TF 0 versus TF 3–10%, 10–20% and above 20%. **(F)** Tumor fraction concordance: predicted versus reference TF in held-out samples with TF ≥3% (97 patients); *r*, Pearson correlation; *P*, two-sided permutation test exchanging patients as whole blocks (**Methods**); dashed line, equality. **(G)** The 50-MAP model and pretrained Fragle^43^ on the same held-out samples of the capture batch: AUC for TF 0 versus ≥3%, balanced accuracy at the tuning threshold, mean absolute error (percentage points, pp) and Pearson r in samples with TF ≥3%. Fragle was run unchanged on the reads outside a target mask inferred from coverage, with its threshold and calibration taken from the same tuning patients (**Methods**). Bars, estimates with 95% patient-bootstrap confidence intervals (1,000 resamples); *, the paired 95% confidence interval of the MAPs-minus-Fragle difference excludes zero. **(H)** Foundation Medicine breast (336 patients; 218 with TF 0 and 118 with TF ≥3%) and lung cancer (322 patients; 260 and 62): ROC curves for TF 0 versus ≥3% for the three models of C, refit within each cancer by nested cross-validation; AUCs are listed in each plot. **(I)** MAP-predicted versus reported TF in patients with TF ≥3% (breast, 118; lung, 62); *r*, Pearson correlation; *P*, two-sided *t*-test. In D–G, each patient is weighted equally. AUC, area under the ROC curve; ROC, receiver operating characteristic.

We next tested whether MAPs represented on existing gene panels could estimate tumor fraction. In 59 patients of the Foundation Medicine cohort with advanced prostate cancer who underwent FoundationOne cfDNA sequencing at Dana-Farber (**Figure S9A**), a model using the 1,342 MAPs that the panel covered with at least ten fragments at both positions in most specimens separated samples with a reported tumor fraction (Foundation Medicine estimate) of 0% from those with ≥3% (AUC = 0.89; Figure 5C), a cutoff close to the detection limit of copy-number-based estimators.^42^ The MAP model outperformed models based on copositioned anchors, sequencing depth, or fragment-length and sequencing features (**Figure 5C**; **Figure S9I**). In 16 test patients, MAP predictions correlated strongly with reported tumor fraction (Spearman ρ = 0.92; **Figure S9B,C**), with ρ = 0.75 and 0.64 in breast and lung cancer (**Figure S9D–G**). The 1,342 MAPs spanned only 0.23 Mb, indicating that tumor fraction-associated nucleosome positioning could be measured from a small targeted genomic footprint.

We further tested this approach in two capture batches from the Virtanen–Kwan study,^55^ whose different panel designs covered 62 and 99 measurable MAPs. In the capture batch, a 50-MAP model trained in 494 patients and selected in 157 tuning patients separated reference tumor fraction of 0% from ≥3% in 163 held-out patients (AUC = 0.91) and outperformed models based on copositioned anchors or sequencing depth (**Figure 5D**). Performance increased with tumor fraction (AUC = 0.58, 0.85 and 0.99 at 3–10%, 10–20% and >20%; **Figure 5E**), and predicted tumor fraction correlated strongly with the reference estimate among samples with tumor fraction ≥3% (*r* = 0.89; **Figure 5E,F**). The independent 50-MAP analysis in the additional capture batch gave similar results (AUC = 0.91; Figure S10).

On the same held-out samples, we compared the 50-MAP model with Fragle, a pretrained deep-learning model that estimates tumor fraction from cfDNA fragment lengths and can be run on the off-target reads of targeted panels.^43^ With Fragle’s threshold and linear calibration taken from the same tuning patients that set the MAP threshold, the 50-MAP model separated reference tumor fraction 0 from ≥3% better than Fragle (AUC = 0.91 versus 0.81) and estimated tumor fraction more accurately (mean absolute error, 8.2 versus 13.7 percentage points; **Figure 5G**).

Because the MAPs that track tumor DNA and the direction of their shift depend on the lineages contributing to plasma,^56^ we refit the weights of MAP models within patients of other cancer types, using the same panel MAPs and no new ChIP data. In Foundation Medicine patients with breast (336 patients) or lung cancer (322 patients), MAP models refit by nested cross-validation separated reported tumor fraction 0 from ≥3% (AUCs = 0.93 and 0.96) better than copositioned anchors (AUCs = 0.81 and 0.82) or sequencing depth (AUCs = 0.50 and 0.50; **Figure 5H**; **Figure S9I**), and their estimates correlated with reported tumor fraction (*r* = 0.83 and 0.78; **Figure 5I**). Models transferred from the Virtanen–Kwan cohort, ridge regressions and fragment-length models gave the same conclusions (Figure S11; Figure S12A–C). In a fixed split of the same patients, MAP models separated tumor fraction 0 from ≥3% better than Fragle in 70 breast and 66 lung cancer test patients (AUCs = 0.94 versus 0.80 and 0.96 versus 0.85), with similar errors (7.5 versus 8.8 and 12.3 versus 12.5 percentage points; **Figure S12D,E**).

These results show that tumor fraction-associated MAP signals persist after hybrid capture and can estimate tumor fraction from targeted panels covering a small genomic footprint.

### MAP estimates track changes in tumor fraction within patients

Because changes in tumor fraction can associate tightly with treatment response and tumor progression,^57^ we next asked whether MAP-based estimates tracked changes in tumor fraction within individual patients. Across 69 pairs of consecutive plasma draws (two samples taken one after the other from the same patient) from 49 held-out patients of the Virtanen–Kwan capture batch, changes in MAP-predicted tumor fraction closely correlated with changes in reference tumor fraction (*r* = 0.91), compared with *r* = 0.21 for copositioned anchors (**Figure 6A**). When reference tumor fraction changed by at least 5 percentage points, MAP estimates changed in the same direction in 94% of pairs, compared with 57% for copositioned anchors and 60% for sequencing depth (**Figure 6B**), and they tracked changes between serial draws more closely than Fragle (*r* = 0.91 versus 0.58; **Figure 6C**). This relationship persisted across successive draws in patients with at least three serial samples (**Figure 6D,E**; **Figure S13**). These results show that MAP-based scores from cfDNA follow changes in tumor fraction over time within patients.

**Figure 6.**
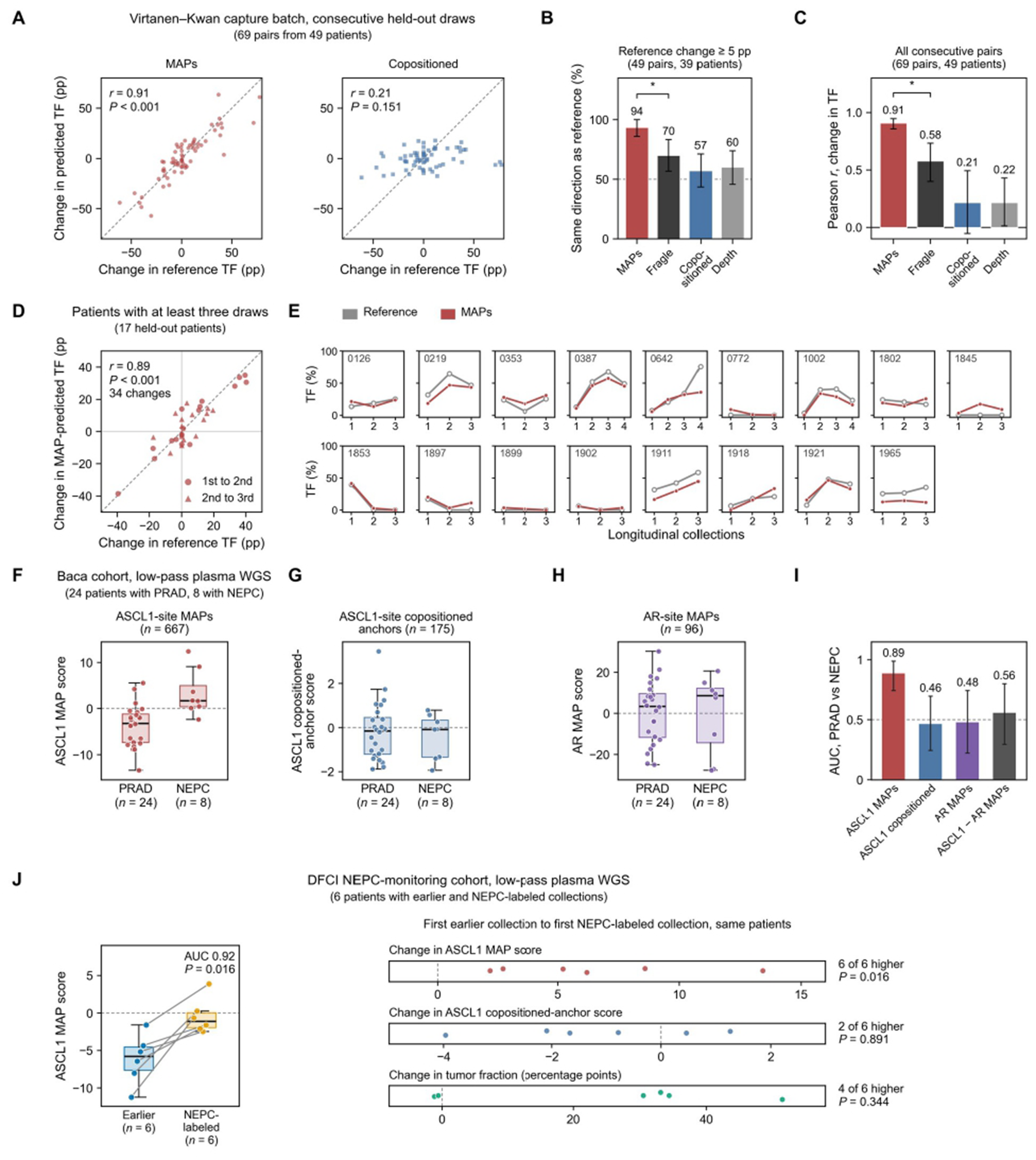
MAPs follow tumor fraction and neuroendocrine state over time. (A) Change in tumor fraction concordance in the Virtanen–Kwan capture batch: change in predicted versus reference tumor fraction (TF) between consecutive draws of the same held-out patient for the 50-MAP model of Figure 5D (left; 69 pairs from 49 patients) and copositioned anchors (right; 65 pairs from 48 patients with copositioned predictions); *r*, patient-weighted Pearson correlation; *P*, two-sided permutation test exchanging patients as whole blocks; dashed line, equality; pp, percentage points. **(B)** Percentage of pairs in which the predicted change had the same sign as the reference change, among pairs with a reference change of at least 5 percentage points (49 pairs from 39 patients; 46 pairs for copositioned anchors) for the four models of C; bars and * as in C; dashed line, 50%. **(C)** The 50-MAP model and pretrained Fragle^43^ on the same serial pairs, with copositioned anchors and sequencing depth: Pearson *r* for changes in TF between consecutive draws (69 pairs from 49 patients; 65 for copositioned anchors); bars, estimates with 95% patient-bootstrap confidence intervals (1,000 resamples); *, the paired 95% confidence interval of the MAPs-minus-Fragle difference excludes zero; Fragle run as in Figure 5G. **(D) Change in MAP-predicted TF against change in reference TF (the source study’s ctDNA fraction) from the first** to the second (circles) and from the second to the third draw (triangles) in held-out patients with at least three draws (34 changes from 17 patients); *r* and *P* as in A; dashed line, equality. **(E)** Reference TF (gray) and MAP-predicted TF (red) at each longitudinal collection for the patients in D (patient numbers of the source study). **(F–H)** Baca cohort, low-pass plasma WGS:^20^ difference between the H3K4me2-and H3K9me3-associated positions in WPS per fragment, averaged per patient, for prostate adenocarcinoma (PRAD; *n* = 24) and neuroendocrine prostate cancer (NEPC; *n* = 8) at 667 MAPs overlapping ASCL1 binding sites in small-cell lung cancer lines (F), at 175 copositioned anchors overlapping the same sites (G) and at 96 MAPs overlapping androgen receptor (AR) binding sites recurrent in primary prostate tumors (H). **(I)** AUC for PRAD versus NEPC, with patient-stratified bootstrap 95% confidence intervals (20,000 draws), for the scores in F–H and for the prespecified ASCL1-minus-AR MAP score (**Methods**). Restricting F to the 220 MAPs at ASCL1 peaks present in all four prostate NEPC ASCL1 ChIP-seq profiles gave an AUC of 0.76 (0.55–0.94). **(J)** DFCI NEPC-monitoring cohort, low-pass WGS: left, ASCL1 MAP score in earlier collections (labeled adenocarcinoma or pre-NEPC) and in NEPC-labeled collections, median per patient and state, in the six patients with both (lines join each patient’s two values); AUC and one-sided exact within-patient permutation *P*. Right, change from the first earlier collection to the first NEPC-labeled collection in the same six patients for the ASCL1 MAP score, the ASCL1 copositioned-anchor score and reported tumor fraction; *P*, one-sided exact sign test. Box plots show the median, interquartile range (IQR) and whiskers to the most extreme values within 1.5 × IQR. In A–D, each patient is weighted equally.

### ASCL1-site MAP protection distinguishes adenocarcinoma and neuroendocrine states of prostate cancer

We next asked whether MAP protection could capture tumor state in addition to tumor fraction by focusing on MAPs overlapping lineage transcription-factor binding sites. In low-pass plasma WGS from the Baca cohort,^20^ protection at 667 MAPs overlapping ASCL1 binding sites distinguished patients with neuroendocrine prostate cancer (NEPC) from those with prostate adenocarcinoma (AUC = 0.89; 95% CI, 0.74–0.99), whereas copositioned anchors overlapping the same ASCL1 binding sites did not (AUC = 0.46; Figure 6F–I).

We then examined serial plasma samples from nine patients monitored for neuroendocrine transformation in the DFCI NEPC-monitoring cohort. Among the six patients with samples before and after an NEPC-labeled timepoint, ASCL1 MAP protection increased in every patient, whereas the corresponding copositioned-anchor signal increased in only two (**Figure 6J**). These results provide an initial example of how cfDNA protection at MAPs at lineage-factor sites can distinguish cancer states.

## DISCUSSION

This study describes loci that we term MAPs (modification-associated positions), where nucleosomes carrying H3K4me2 and H3K9me3 occupy reproducibly offset, partly overlapping positions. By measuring histone modifications associated with different chromatin states from the same plasma sample, our approach controls for variation in contributing cell types and other sample-specific factors, enabling us to discover 35,120 high-confidence MAPs from a relatively small cohort of prostate cancer plasma samples. MAPs were reproducible in an independent cfChIP-seq cohort, its inter-laboratory replicates and MNase-digested prostate cancer cell chromatin across several histone modifications associated with active and inactive chromatin. While prior fragmentomic studies have inferred chromatin state from fragment lengths, end motifs, and coverage over regions,^1,4,5,23^ MAPs tie histone modification states to distinct nucleosome positions. Fragment protection at these fixed positions, measured from cfDNA without immunoprecipitation, correlated with tumor fraction in whole-genome and capture sequencing datasets. A model using cfDNA fragments from 50 MAPs (spanning less than one hundred-thousandth of the human genome) estimated tumor fraction in held-out patients and cohorts and tracked changes across serial draws. Models transferred between prostate cancer cohorts on shared MAPs without refitting. Despite using MAPs discovered only in prostate cancer plasma, models fit in breast and lung cancer cohorts separated tumor fractions better than fragment-length and sequencing features. Together, these results establish a compact positional readout of tumor fraction that can be read from targeted panels already used in ctDNA profiling.

MAPs were widely distributed across the genome and differed from copositioned anchors mainly by their depletion near CTCF and cohesin sites and promoters and their enrichment in the bodies of expressed genes and in distal enhancer-like elements. CTCF binding is flanked by regularly phased nucleosome arrays, and CTCF and cohesin sites have distinctive local nucleosome organization.^35,51,52,58^ The depletion at MAPs was strongest for promoter activity shared across the assessed prostate, neutrophil, and monocyte lineages and for CTCF sites bound in both prostate and blood cells. Among MAPs, greater separation was associated with fewer nearby CTCF and cohesin peaks. We propose that sites bound in every contributing cell, by CTCF and cohesin or by RNA polymerase II at promoters, set nucleosomes at similar positions regardless of chromatin state, as at copositioned anchors. Away from such shared barriers, factors that position nucleosomes, such as transcription factors and chromatin remodelers, may act in active chromatin of one cell type but not in repressed chromatin of another, so that nucleosomes carrying the two marks occupy offset positions. The balance between the two positions in plasma would then reflect the chromatin states of the contributing cells (**Figure 3F**).

Because nucleosome protection signals can be recovered from targeted panels,^5,59^ MAPs could be added to assays already used in cancer profiling. Tumor fraction is often estimated from variant allele fractions at a few mutated sites, but variants can arise from clonal hematopoiesis (CHIP) rather than from cancer: 53.2% of cfDNA variants in patients with cancer had features of CHIP,^60^ and 10% of men with advanced prostate cancer carried CHIP-related variants in genes used to select PARP inhibitor treatment.^61^ Copy-number-based estimates are also affected: clonal hematopoiesis can produce copy-number changes in cfDNA that mimic tumor DNA.^62,63^ MAP estimates use neither mutations nor copy-number changes but fragment positions at loci whose chromatin differs between tumor and blood cells, so testing them in patients with known CHIP is a direct next step. MAPs could provide an estimate when no tumor-derived coding alteration is detected, as was reported in 15% of 21,807 patients with advanced cancer,^64^ and could complement current methods that require assumptions about clonality, heterozygosity, and copy number. Other fragmentomic-based approaches have been reported for estimating tumor fraction.^43,59^ A potential advantage of MAPs is that cfDNA-inferred nucleosome positions are robust across datasets with variation in library construction and hybrid-capture approaches. In contrast, the large degree of technical variation in cfDNA fragment lengths between samples may pose challenges for methods that use fragment lengths as inputs.

Our study has several limitations. MAPs were derived from bulk profiles and separate immunoprecipitations of the same sample, so we cannot establish co-occupancy or movement of a single nucleosome. Some copositioned anchors may also reflect background fragments recovered by both immunoprecipitations at highly accessible promoters, rather than two marked nucleosomes at the same position.^65,66^ The separation cutoffs follow the helical turn of DNA and the size of the nucleosome core, but these and other choices, such as the 30-bp cross-cohort matching tolerance, affect the set of MAPs called; matching tolerances of 20 and 50 bp, for example, gave 26,810 and 45,605 MAPs (**Figure S1E,G–I**; **Supplementary Note 1**). MAPs were discovered in plasma from patients with prostate cancer, and studies in other cancers will be needed to establish how widely they occur. The WGS controls came partly from a different study, the capture comparison group was relatively small, and reference tumor fractions came from different estimators in different cohorts. Both capture held-out sets came from one source cohort. The lineage analyses are exploratory: the prespecified ASCL1-minus-AR score did not distinguish PRAD from NEPC. Prospective cohorts and further comparisons with existing approaches will be needed to assess the utility of MAPs for estimating and monitoring tumor fraction.

Our results indicate that the genomic positions of circulating nucleosomes retain a measurable trace of the histone modifications of the cells that released them. Because MAP positions can be added to targeted panels already used in cancer care, MAP protection could serve as an additional feature for tumor detection and for monitoring response without a separate assay. MAPs at lineage transcription-factor sites already distinguished some neuroendocrine states (**Figure 6F–J**). Discovered in other cancers and cell types, and combined with single-nucleosome epigenetic assays,^67^ MAPs could make the regulatory programs of tumors readable from routine cfDNA sequencing.

## ACKNOWLEDGEMENTS

We thank members of the Freedman and Baca laboratories for discussions and feedback. We are grateful to the patients who contributed plasma to the studies analyzed here, and to the investigators of Sipola et al., Herberts et al. and Virtanen, Kwan et al. for making their data available. We thank Foundation Medicine for providing de-identified assay data. This work was supported by the Fund for Innovation in Cancer Informatics, the Wong Family, the Damon Runyon Cancer Research Foundation, the US Department of Defense (DoD) (award W81XWH-21-1-0358), the Prostate Cancer Research Foundation, the National Institutes of Health (NIH) (1U01CA296432-01), and a gift from Jeffery Parker. G.S.G. is supported by the National Cancer Institute (T32CA009172) and the Lubin Family Foundation Scholar Award. This research was supported in part by the Intramural Research Program of the National Institutes of Health (NIH) (ZIABC011965 to C.F. Tyndall and D.Y. Takeda, ZIABC011680 and ZIABC011838 to A.G.S.). The contributions of the NIH authors are considered Works of the United States Government. The findings and conclusions presented in this paper are those of the authors and do not necessarily reflect the views of the NIH or the U.S. Department of Health and Human Services. Next-generation sequencing was done at the CCR Genomics Core at the National Cancer Institute, NIH, Bethesda, MD. Figures were created in part with BioRender.com.

## AUTHOR CONTRIBUTIONS

Conceptualization, Y.O., G.S.G., D.Y.T., and S.C.B.; Methodology, Y.O., G.S.G., and S.C.B.; Software and Formal Analysis, Y.O., G.S.G., Ze.Z., M.P., Z.J., Zi.Z. and E.I.; Generation and analysis of MNase-ChIP and cfDNA sequencing datasets, J.-H.S., Ze.Z., G.G.L., T.A.H., S.D.S., J.L., A.G.S., R.N., G.N., J.E.B., F.K. and D.Y.T.; Generation of cfChIP-seq data, C.F.T. and D.Y.T.; Data Curation, Y.O., G.S.G., J.-H.S., Ze.Z., M.P. and Z.J.; Writing – Original Draft, Y.O.; Review & Editing, all authors; Visualization, Y.O.; Supervision, G.S.G., S.C.B. and D.Y.T.; Funding Acquisition, S.C.B.

## SUPPLEMENTARY FIGURES

**Figure S1.**
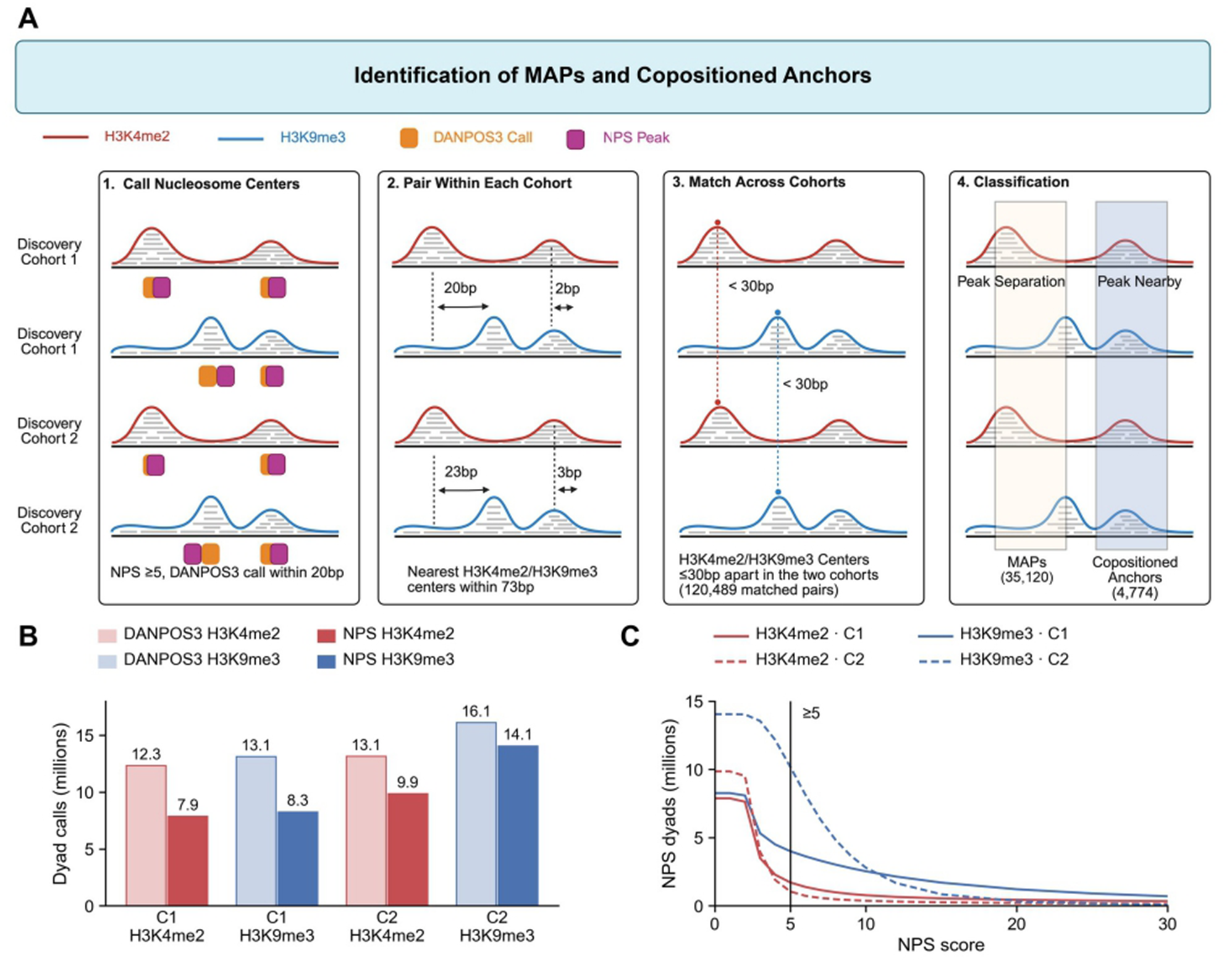
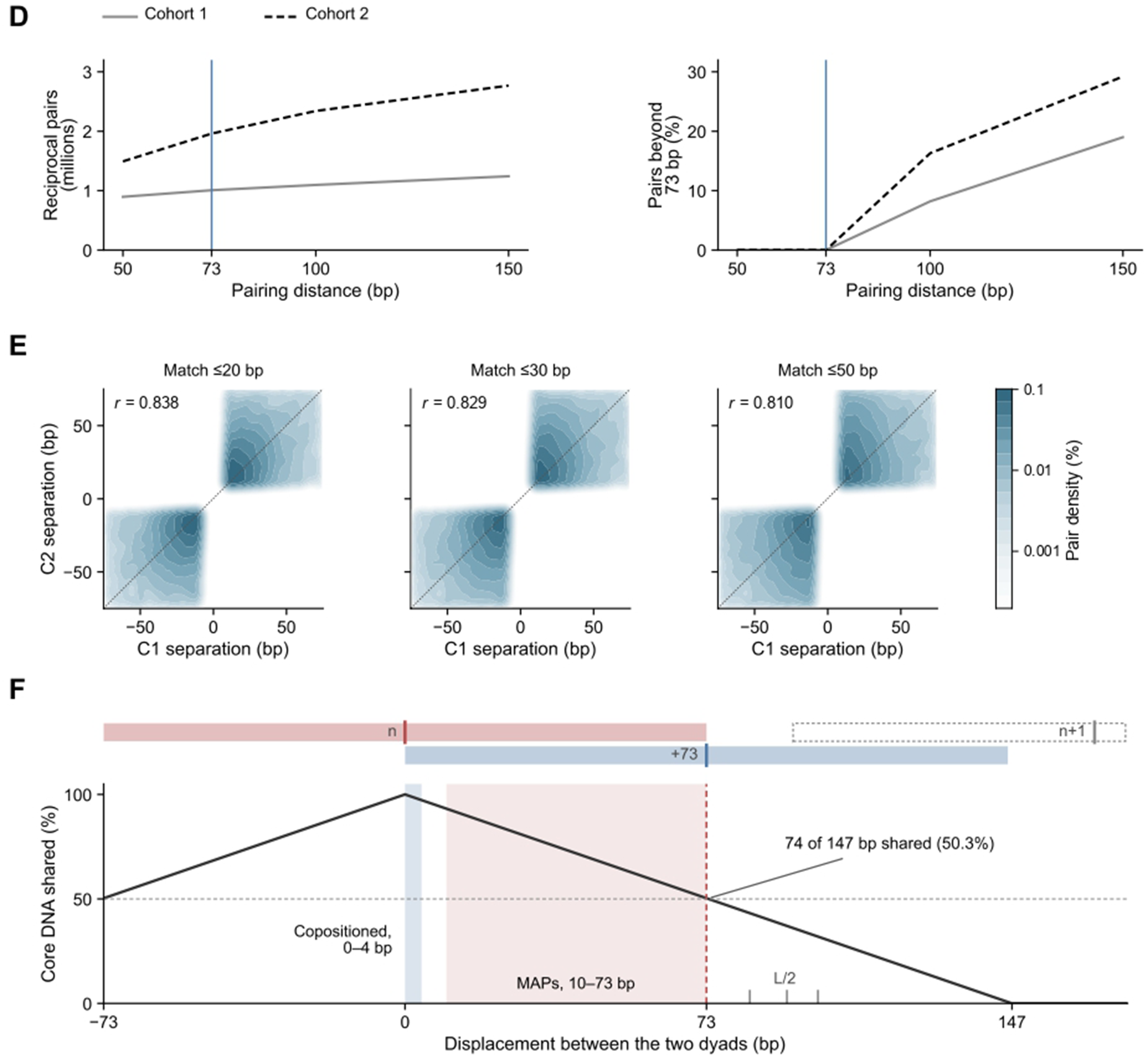
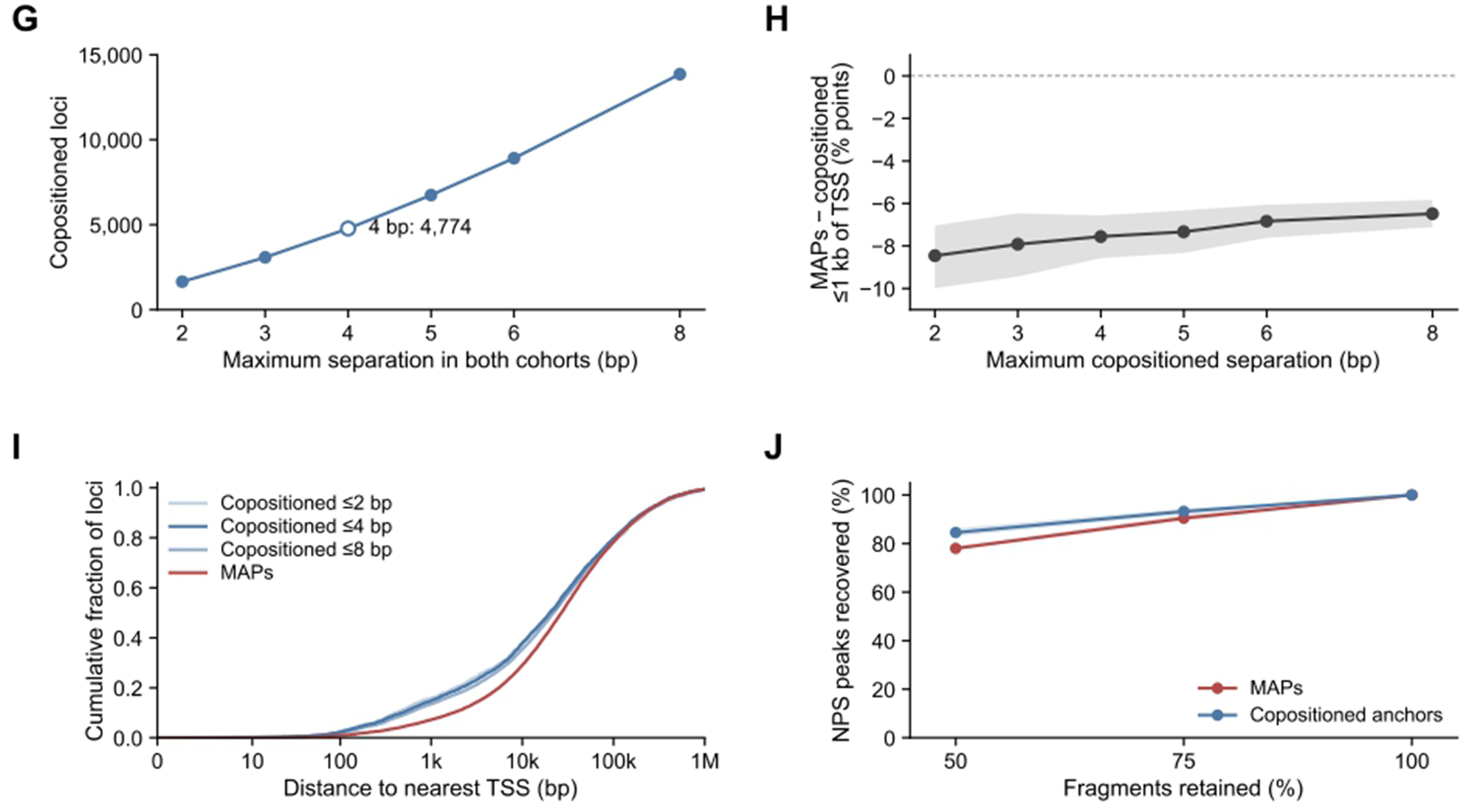
Identification of MAPs and copositioned anchors, and robustness of the calls to cutoffs and read depth. **(A)** Discovery of MAPs (histone modification-associated positions) across the two discovery cohorts (cohort 1, 7 participants and 17 specimens; cohort 2, 8 participants and 20 specimens, 19 with both marks). High-confidence nucleosome positions were defined as dyads with a probabilistic nucleosome score (PNS) ≥5 and a DANPOS3 dyad call within 20 bp; the PNS peak center was kept as the dyad position. H3K4me2- and H3K9me3-associated dyads were paired within 73 bp in each cohort. Pairs were then matched between the two cohorts when their H3K4me2-associated dyads were reciprocal nearest neighbors within 30 bp. MAPs are matched pairs with the same order of the two marks in both cohorts and a separation of 10–73 bp in both cohorts; copositioned anchors are separated by 0–4 bp in both cohorts. The resulting atlas comprises 120,489 paired loci: 35,120 modification-associated position (MAP) loci, 4,774 copositioned H3K4me2/H3K9me3 anchors, and 80,595 loci meeting neither definition. **(B)** DANPOS3 (light) and PNS (dark) dyad-call counts in each cohort (C1, C2) and mark. **(C)** PNS dyads at or above each score threshold; the vertical line marks the threshold used (≥5). Colors identify histone marks; solid and dashed lines identify discovery cohorts. **(D)** Number of reciprocal nearest-neighbor pairs (left) and percentage of pairs separated by more than 73 bp (right) as the maximum pairing distance increases; vertical lines mark the 73-bp pairing distance used. **(E)** Dyad spacing (signed H3K4me2–H3K9me3 separation) in cohort 1 (C1) against cohort 2 (C2) at cross-cohort matching tolerances of 20, 30 and 50 bp, with the within-cohort H3K4me2–H3K9me3 pairing distance fixed at 73 bp. The corresponding MAP counts are 26,810, 35,120 and 45,605. Colors show pair density on a shared logarithmic percentage scale; dotted diagonals mark equal separation; r is the Pearson correlation. **(F)** Core DNA shared by two dyads as a function of their separation; dyads within 73 bp, half of the 147-bp nucleosome core, share at least 74 bp. Shading marks the separations of copositioned anchors (0–4 bp) and MAPs (10–73 bp). Top, the footprint of nucleosome n (red), a footprint displaced by 73 bp (blue) and the neighboring nucleosome n+1 (dashed). L/2, half the nucleosome repeat length L, is the midpoint between adjacent dyads; ticks mark L/2 for L = 167, 185 and 200 bp (83.5, 92.5 and 100 bp), beyond which a dyad lies nearer the neighboring nucleosome than nucleosome n (**Supplementary Note 1**). **(G)** Number of copositioned loci at each maximum separation required in both discovery cohorts. The open point marks the 4-bp cutoff used (4,774 loci). **(H)** Difference between MAPs and copositioned anchors in the percentage of loci within 1 kb of a transcription start site (TSS) at various distance thresholds for copositioning; negative values mean lower promoter proximity at MAPs. Shading shows pointwise 95% percentile intervals from 5,000 bootstrap samples of the 23 chromosomes, resampling the same chromosomes for both groups. **(I)** Cumulative distributions of nearest-TSS distance for the 2-, 4- and 8-bp copositioned definitions and for MAPs, measured from locus midpoints with hg19 knownGene transcripts. All comparisons in H and I keep the same 35,120 MAPs. **(J)** Recovery of a PNS peak after downsampling fragments to 50%, 75% and 100% at 460 loci chosen without reference to signal (ten MAPs and ten copositioned loci from each of 23 chromosomes; three random seeds per reduced depth). A peak counts as recovered when a positive PNS peak lies within 20 bp of its full-depth position, among profiles with a full-depth peak within 20 bp of the atlas coordinate, the midpoint of the two paired dyads (230 loci per class in both marks and both discovery cohorts, 920 profiles per class; 868 of 920 MAP and 892 of 920 copositioned profiles). Lines show the mean across seeds. These subsampling checks assess PNS peak repeatability at discovery-defined loci.

**Figure S2.**
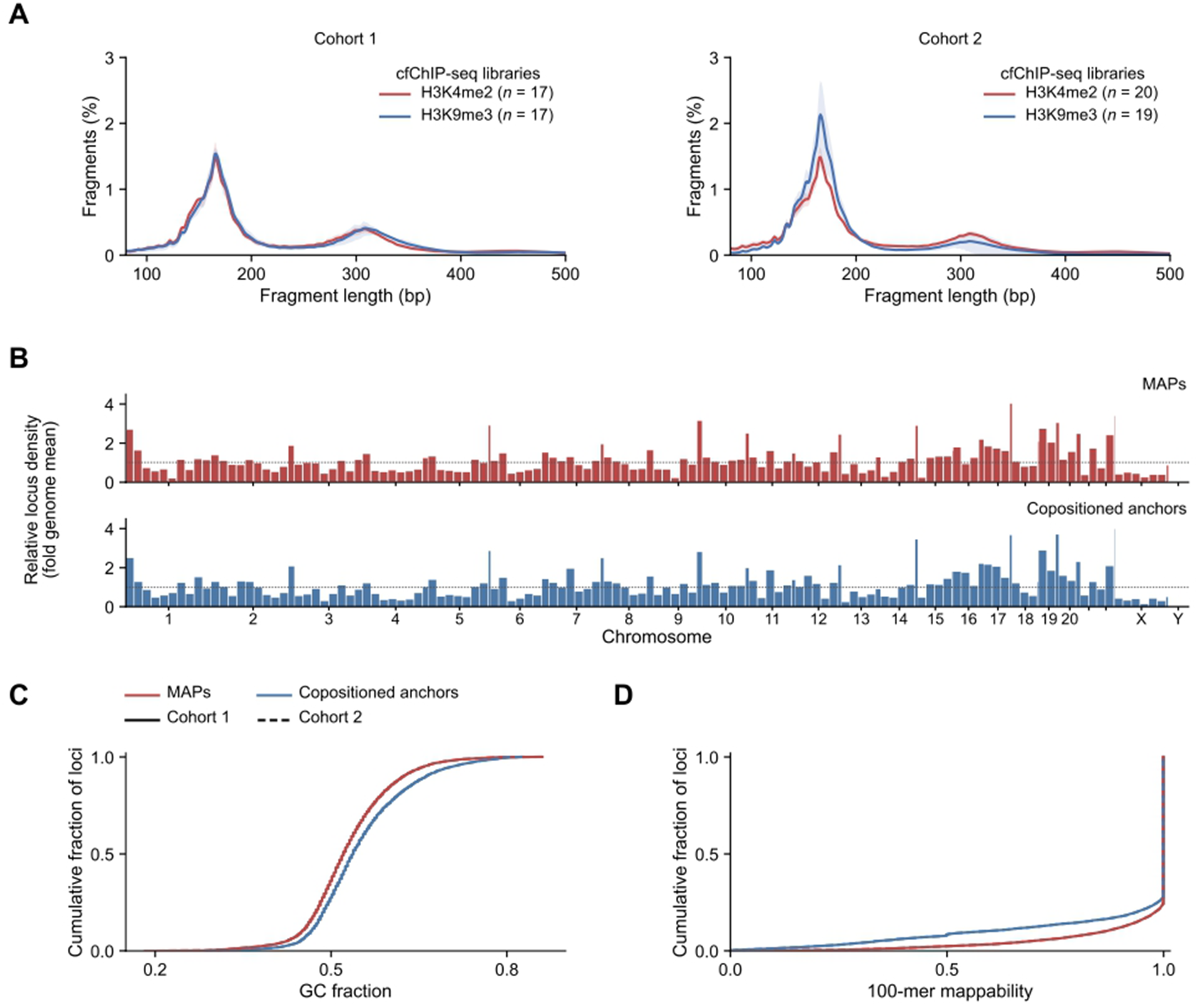
Discovery cfChIP-seq libraries and genomic characteristics of the MAP atlas. **(A)** Fragment-length distributions of the discovery cfChIP-seq libraries. Curves show the mean frequency after normalizing each library over 80–500 bp; shading shows the interquartile range across libraries. Cohort 1 has 17 libraries for each mark; cohort 2 has 20 H3K4me2 and 19 H3K9me3 libraries. **(B)** Locus density in 25-Mb genomic blocks relative to the genome-wide mean of each class (35,120 MAPs, top; 4,774 copositioned anchors, bottom); the dotted line marks one. **(C, D)** Cumulative distributions of GC fraction **(C)** and 100-mer mappability **(D)** in 301-bp windows centered on the paired-position midpoints; solid and dashed curves represent cohorts 1 and 2, respectively. Median GC content was 52.2% at MAPs and 53.8% at copositioned anchors. In B–D, red denotes MAPs and blue denotes copositioned anchors.

**Figure S3.**
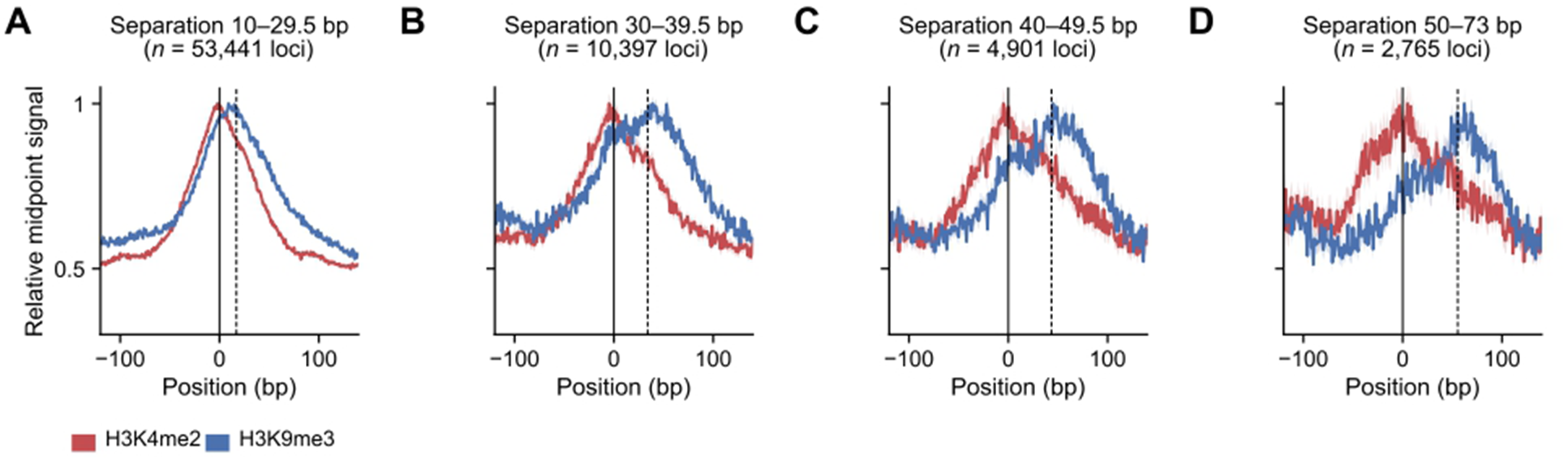
Independent-cohort cfChIP-seq profiles at discovery-paired loci, grouped by discovery separation. (A–D) Pooled H3K4me2 (red) and H3K9me3 (blue) fragment-midpoint profiles in the independent cohort (Sipola et al., 2025) at 71,504 loci (all 35,120 MAPs and 36,384 other paired loci), grouped by their mean separation in the two discovery cohorts: 10–29.5 bp (A; 53,441 loci), 30–39.5 bp (B; 10,397), 40–49.5 bp (C; 4,901) and 50–73 bp (D; 2,765). Positions, group membership and orientation were fixed from the discovery cohorts. Profiles are centered on the discovery H3K4me2-associated position (solid line), with loci whose H3K9me3 position lies upstream reversed to a common orientation; dashed lines mark the median discovery separation in each group (17, 34, 43.5 and 55.5 bp). Curves are mean midpoint signal across loci, normalized separately for each mark and group to the maximum over ±250 bp and shown from −120 to +140 bp; shading is the standard error across loci. The pooled profiles test whether the displacement measured in discovery recurs in independent plasma.

**Figure S4.**
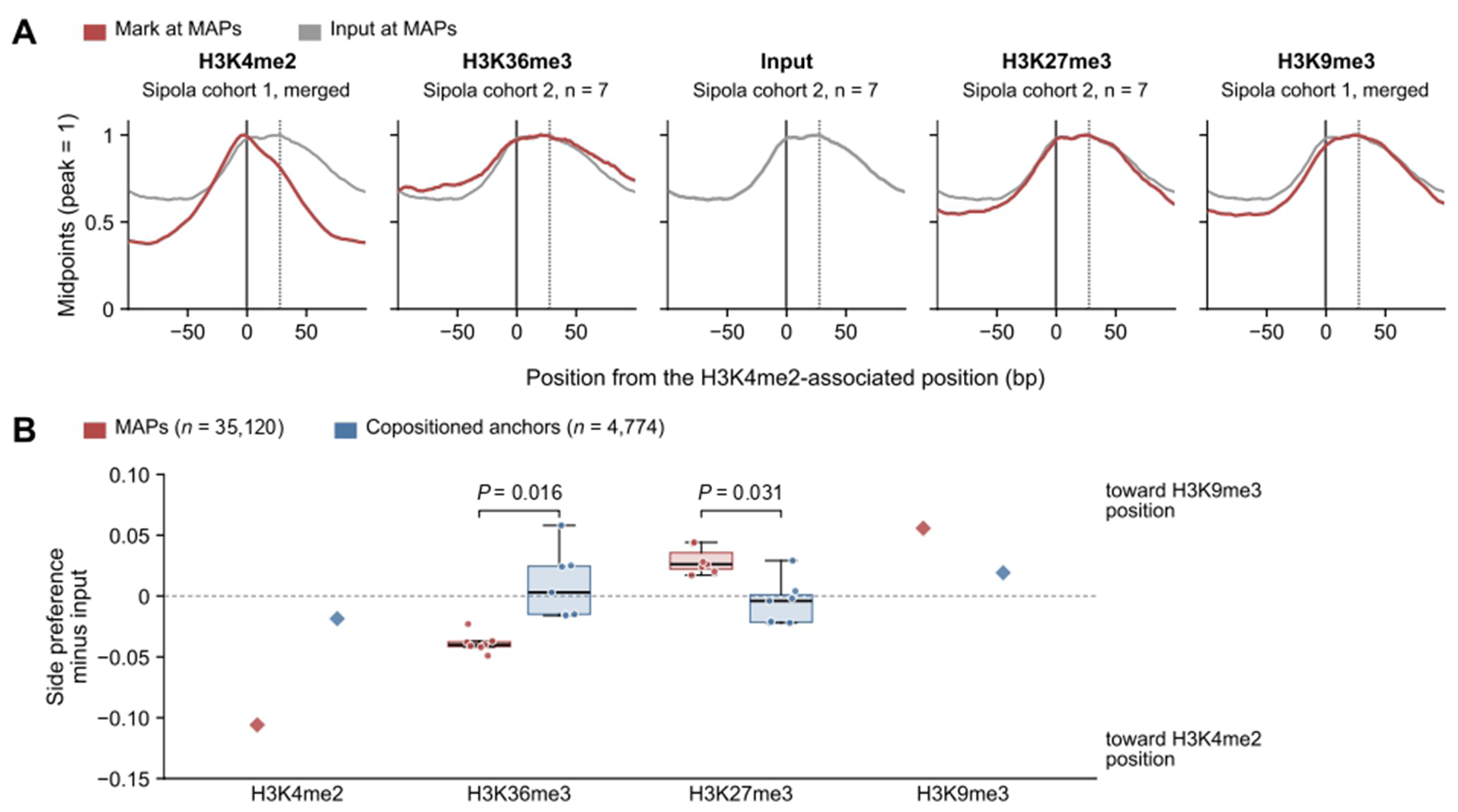
Other histone marks along the MAP axis, each read against input. Fragment midpoints (120–180-bp fragments) at the 35,120 MAPs and the 4,774 copositioned anchors. Zero is the H3K4me2-associated position and positive positions point toward the H3K9me3-associated position. Libraries: H3K4me2 and H3K9me3, the merged independent-cohort libraries of Figure 2 (Sipola et al., 2025); H3K27me3, H3K36me3 and input, the seven inter-laboratory replicate specimens, each with H3K27me3, H3K36me3 and input libraries from the same plasma. Profiles of H3K27me3, H3K36me3 and input are summed over their libraries. **(A)** Midpoint profile of each mark at MAPs (red) over the input profile at the same MAPs (gray), each scaled to its peak and smoothed over 11 bp. The dotted line marks the median H3K9me3-associated position (+28 bp). Input fragments are also displaced at MAPs. **(B)** Side preference of each mark minus that of input, at MAPs (red) and at copositioned anchors (blue). Side preference is (R − L)/(R + L), where R and L are fragment midpoints at +10 to +73 bp and −73 to −10 bp; positive values lie on the H3K9me3 side of input. H3K27me3 and H3K36me3: each specimen minus its own input (7 specimens each); H3K27me3 lies further toward the H3K9me3 position than its input in 7 of 7 specimens (median +0.026) and H3K36me3 further toward the H3K4me2 position in 7 of 7 (median −0.040). H3K4me2 and H3K9me3: merged libraries minus summed inter-laboratory replicate input (one value each). MAPs and copositioned anchors differ in H3K36me3 (7 specimens; *P* = 0.016) and H3K27me3 (7 specimens; *P* = 0.031), by exact two-sided Wilcoxon signed-rank tests of the paired differences. Box plots show the median (center line) and the interquartile range (box); whiskers extend to the most extreme values within 1.5 times the interquartile range.

**Figure S5.**
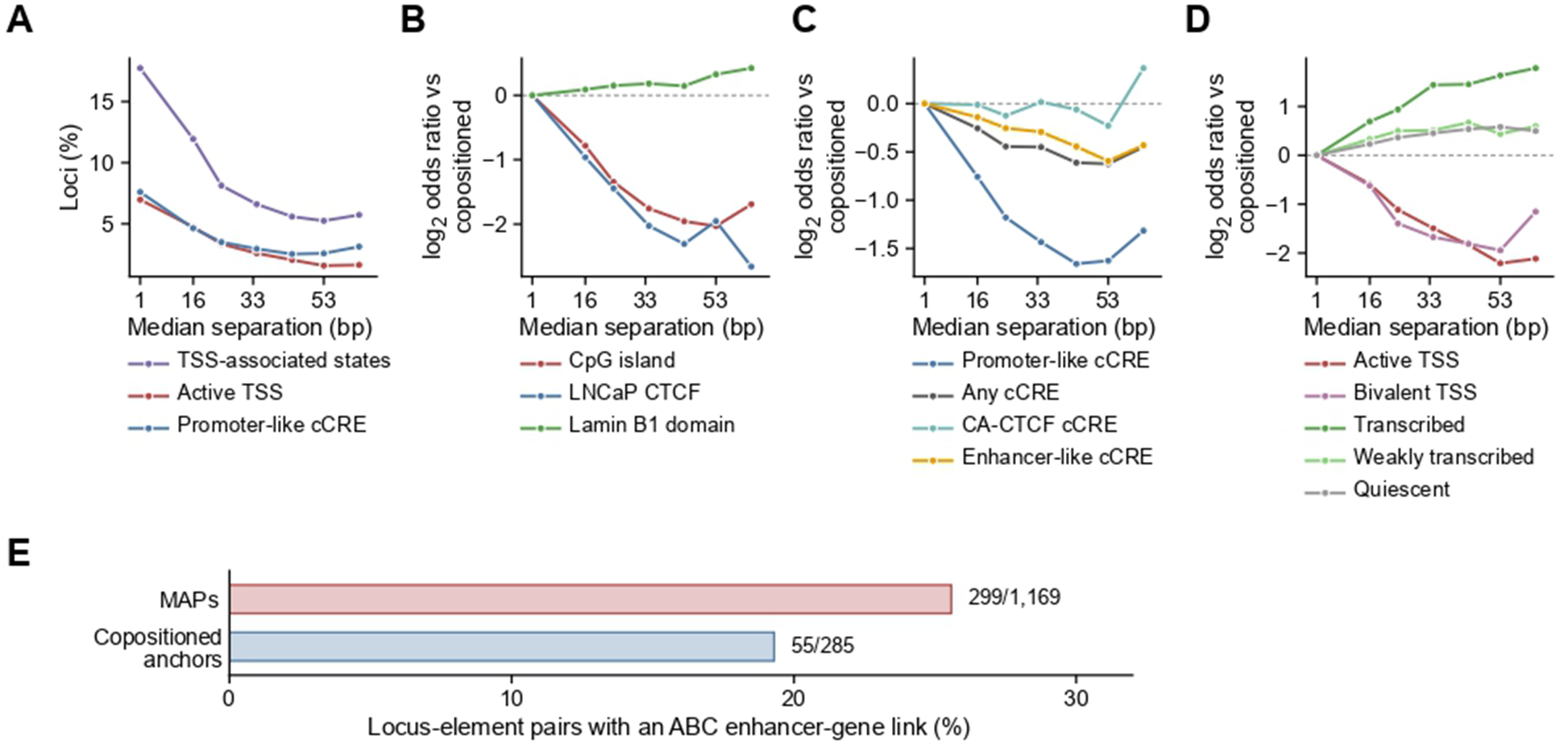
Genomic and regulatory context of MAPs across displacement strata, and LNCaP enhancer–gene links. **(A)** Frequencies of active transcription-start-site (TSS) states, TSS-associated states and promoter-like candidate cis-regulatory elements (cCREs). **(B)** Association with CpG islands, LNCaP CTCF sites and Lamin B1-associated domains. **(C)** Composition of SCREEN v4 cCRE classes. **(D)** Prostate EpiMap chromatin states. In A–D the complete copositioned anchor class (*n* = 4,774; median separation 1 bp) is compared with MAPs in six displacement strata, 10–19, 20–29, 30–39, 40–49, 50–59 and 60–73 bp (*n* = 7,629, 11,352, 8,334, 4,976, 2,096 and 733 loci), which together comprise all 35,120 MAPs; points sit at the median separation of each stratum (16, 24, 34, 44, 53 and 63 bp). A shows percentages; B–D show log2 odds ratios relative to the complete copositioned class, with a 0.5-cell correction. Binary annotations use 151-bp midpoint windows and chromatin states are assigned at the midpoint. TSS-associated states combine active, flanking and bivalent TSS states; enhancer-like cCREs combine proximal and distal enhancer-like signatures. Lines connect the stratum summaries. **(E)** Percentage of non-promoter LNCaP links between a locus and a regulatory element that predict a distal enhancer–gene contact (Activity-by-Contact [ABC] score ≥0.022): 299 of 1,169 for MAPs and 55 of 285 for copositioned anchors.

**Figure S6.**
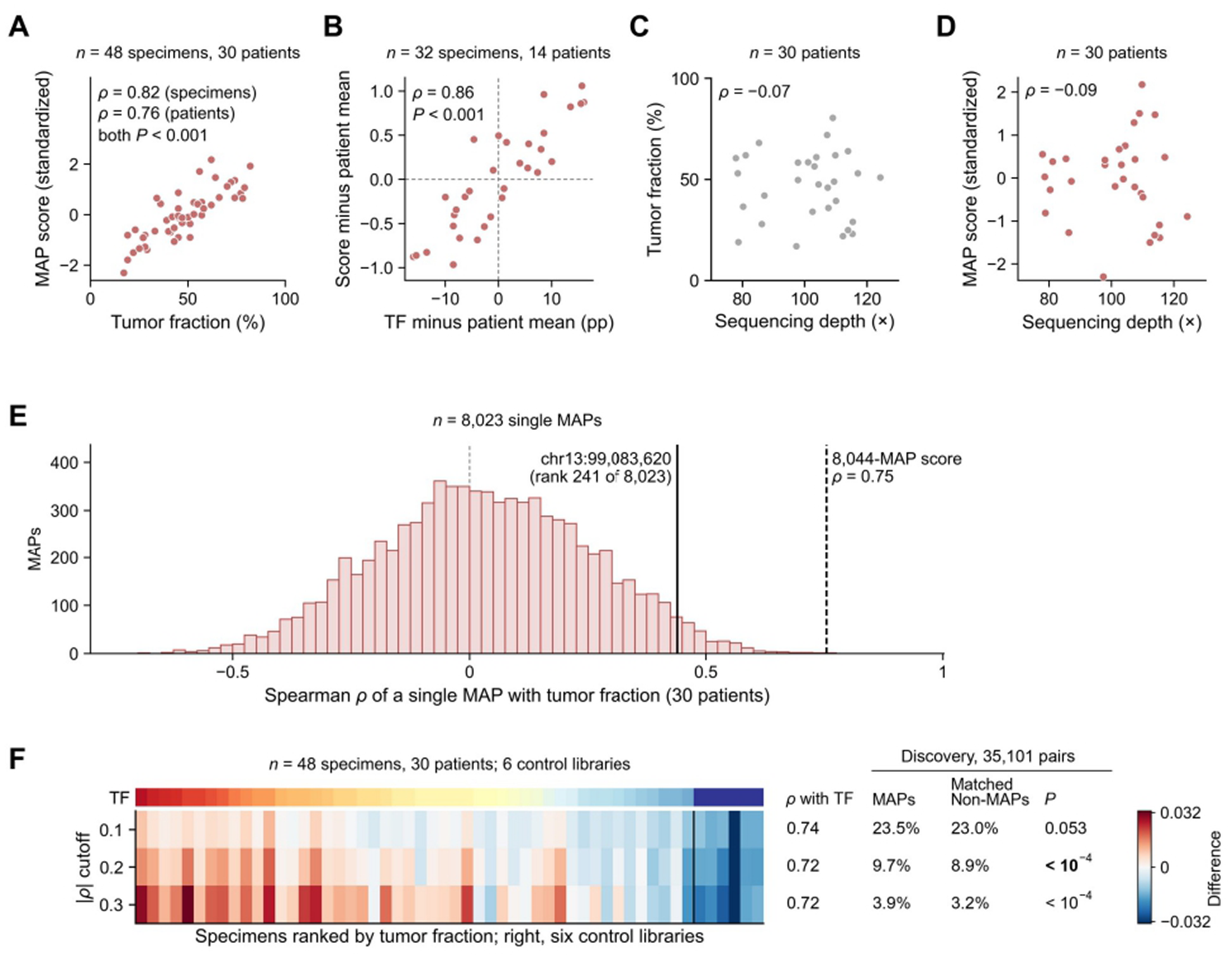
Deep plasma WGS: the MAP score against tumor fraction across and within patients, sequencing depth, single MAPs and the threshold that defines the tumor fraction-correlated groups. Deep plasma WGS from Herberts et al. (Nature, 2022): 48 specimens from 30 patients with metastatic castration-resistant prostate cancer, with tumor fraction (TF) as reported by the source study, and six control libraries assigned TF = 0: four from Snyder et al. (Cell, 2016; BH01, pooled plasma, and IH01–IH03) and two local controls (CTRL-AE-179 and CTRL-AE-201). The score is the standardized mean oriented contrast at 8,044 oriented MAPs (**Methods**). **(A)** Score against reported TF for the 48 cancer specimens; ρ is the Spearman correlation across specimens and across the 30 patient means (95% CI for the patient means from resampling whole patients, 0.48–0.90; permutation P from 10,000 permutations). **(B)** Within patients: the score against TF in the 14 patients with two or three specimens (32 specimens). Both axes are centered within each patient (specimen value minus the patient mean), so only change within a patient is shown; TF is in percentage points (pp). ρ = 0.86 (95% CI from resampling whole patients, 0.64–0.95); permutation *P* = 0.0006 (10,000 permutations). **(C, D)** Estimated sequencing depth against TF **(C)** and against the score **(D)**, one point per patient (30 patients; ρ = −0.07 and −0.09); adjusted for sequencing depth and median fragment size, the patient-level correlation of score and TF was partial ρ = 0.72. **(E)** Spearman correlation of each of the 8,044 MAPs, taken singly, with patient-mean TF across the 30 patients (median ρ = 0.03; 21 MAPs whose contrast did not vary between patients have no correlation and are not drawn). Solid line, the MAP at chr13:99,083,620 (ρ = 0.44; rank 241 of 8,023); dashed line, the 8,044-MAP score (ρ = 0.75), which exceeds all but one single MAP. **(F)** For each specimen (columns: the 48 specimens ranked by TF, then the six control libraries), WPS at the H3K4me2-associated minus the H3K9me3-associated position averaged over K4/TF-correlated MAPs, minus the same average over K9/TF-correlated MAPs, with the two groups of Figure 4A defined at |ρ| > 0.1, 0.2 and 0.3 (rows; 4,435 and 3,822, 1,776 and 1,619, and 681 and 695 of the 35,120 MAPs). Top, TF. Right: Spearman ρ with TF across the 48 specimens (0.83, 0.79 and 0.75; patient means, 0.74, 0.72 and 0.72; patient-label permutation *P* = 5 × 10⁻⁴ at every threshold). Far right: in the 36 discovery specimens, the percentage of 35,101 MAPs and of their matched Non-MAP loci (matched within chromosome on discovery coverage, GC content and mappability) at which H3K4me2 and H3K9me3 correlated with TF in opposite directions beyond each threshold; *P*, exact McNemar test on the pairs. Figure 4 uses |ρ| > 0.2, at which MAPs exceed matched Non-MAP loci (*P* = 6 × 10⁻⁵; at |ρ| > 0.1, *P* = 0.05) and each group keeps more than 1,000 MAPs.

**Figure S7.**
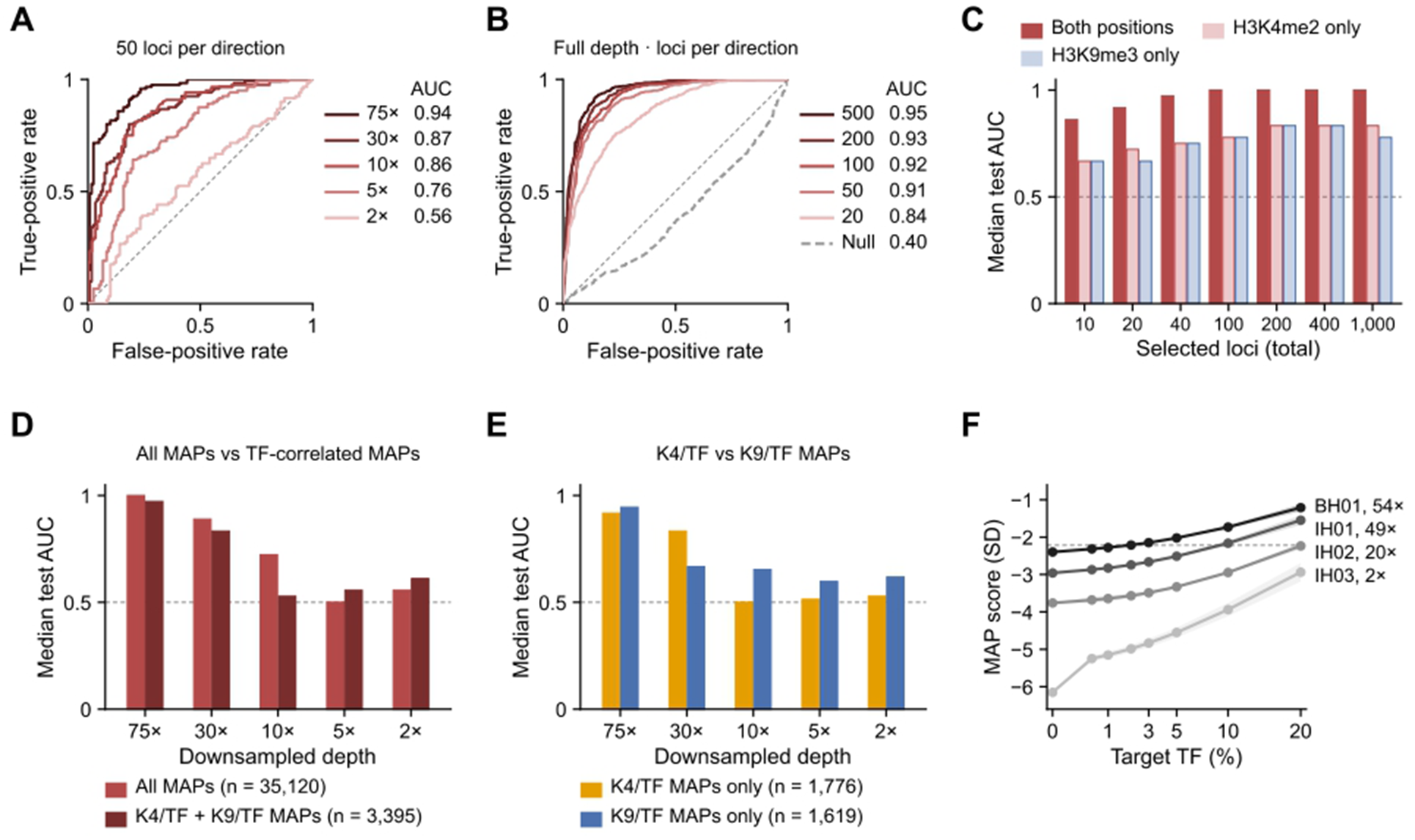
Deep plasma WGS: tumor-fraction quartile discrimination after read downsampling, with fewer loci and by readout, and the MAP score of in-silico mixtures. Deep plasma WGS from Herberts et al. (Nature, 2022). Specimens in the lowest (Q1; tumor fraction [TF] 17–45%) and highest (Q4; 51–82%) quartiles of reported TF were compared. In each repeated partition, half of the specimens were used to rank loci by their Q4-versus-Q1 area under the receiver operating characteristic curve (AUC) and to fix their direction, and the other half were scored; test scores from all partitions are pooled into one empirical ROC curve per setting. Loci in A–C come from the MAPs in both discovery cohorts. (A) Discrimination after downsampling the WGS reads from 75× to 30×, 10×, 5× and 2×, with 50 loci per direction (20 partitions of 12 test specimens; 31–36 specimens per depth): AUCs = 0.94, 0.87, 0.86, 0.76 and 0.56. **(B)** Discrimination at full depth with 20, 50, 100, 200 and 500 loci per direction (100 partitions of 12 test specimens; 19 specimens per quartile): AUCs = 0.84, 0.91, 0.92, 0.93 and 0.95. The direction-null control takes 500 loci from the pool in which H3K4me2 and H3K9me3 signals in the discovery cohort did not move in opposite directions with TF (AUC = 0.40). **(C)** Discrimination between Q4 and Q1 using paired-position, H3K4me2-position-only and H3K9me3-position-only measurements at the same selected loci, at 10–1,000 selected loci in total. Bars are median test AUCs across 100 specimen partitions (24 training and 24 test specimens; six test specimens per quartile); locus selection and direction were set in training data. **(D, E)** Discrimination of Q4 versus Q1 at downsampled depths from 75× to 2× using all MAPs, the K4/TF-correlated plus K9/TF-correlated MAPs, or each of the two groups alone as inputs (35,097, 3,393, 1,776 and 1,617 input features). Bars are median test AUCs over 100 repeated partitions of 24 specimens from 21 patients (twelve training and twelve test specimens); preprocessing and tuning used training data only. **(F)** MAP scores of in-silico mixtures of two cancer specimens (AE-136-Progression, TF 82%; AE-015-Progression2, TF 77%) into each of the four Snyder libraries (BH01 and IH01–IH03; approximate depth shown), at target TF from 0 to 20%. Mixtures were formed from the raw protection profiles of the two libraries, each divided by its number of 120–180-bp fragments and weighted by the target TF, before normalization and smoothing. Lines show the mean of the two cancer specimens; shading, their range; the dashed line marks the lowest of the 48 cancer specimens. The score is the oriented H3K4me2-minus-H3K9me3 contrast at the 8,044 MAPs carried to hg38, averaged with equal weight over the two orientation sets and standardized by the mean and SD of the 48 cancer specimens.

**Figure S8.**
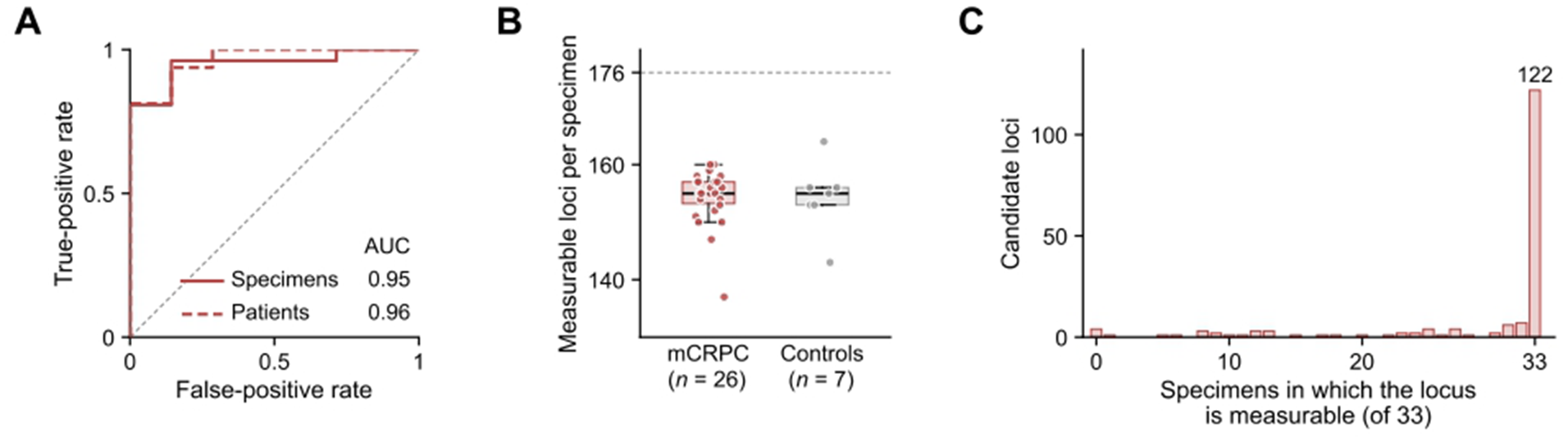
Hybrid-capture sequencing in the Emory capture cohort: mCRPC versus cancer-free controls and measurable loci. Emory capture cohort: 33 plasma specimens from 16 patients with metastatic castration-resistant prostate cancer (mCRPC; 26 specimens) and 7 cancer-free controls (7 specimens). The candidate-locus score contrasts protection at the H3K4me2- and H3K9me3-associated positions across 176 candidate loci, expressed per 100 fragments. A candidate locus is measurable in a specimen when at least one fragment covers each of its two positions. **(A)** ROC curves separating mCRPC from cancer-free controls, scored once per specimen (solid; 26 and 7 specimens; area under the receiver operating characteristic curve [AUC] = 0.95) and once per patient as the median of that patient’s specimens (dashed; 16 and 7 patients; AUC = 0.96). **(B)** Measurable candidate loci per specimen (dashed line, all 176 candidates). Box plots show the median (center line) and the interquartile range (box); whiskers extend to the most extreme values within 1.5 times the interquartile range. **(C)** Number of specimens in which each candidate locus is measurable: 172 loci in at least one specimen and 122 in all 33.

**Figure S9.**
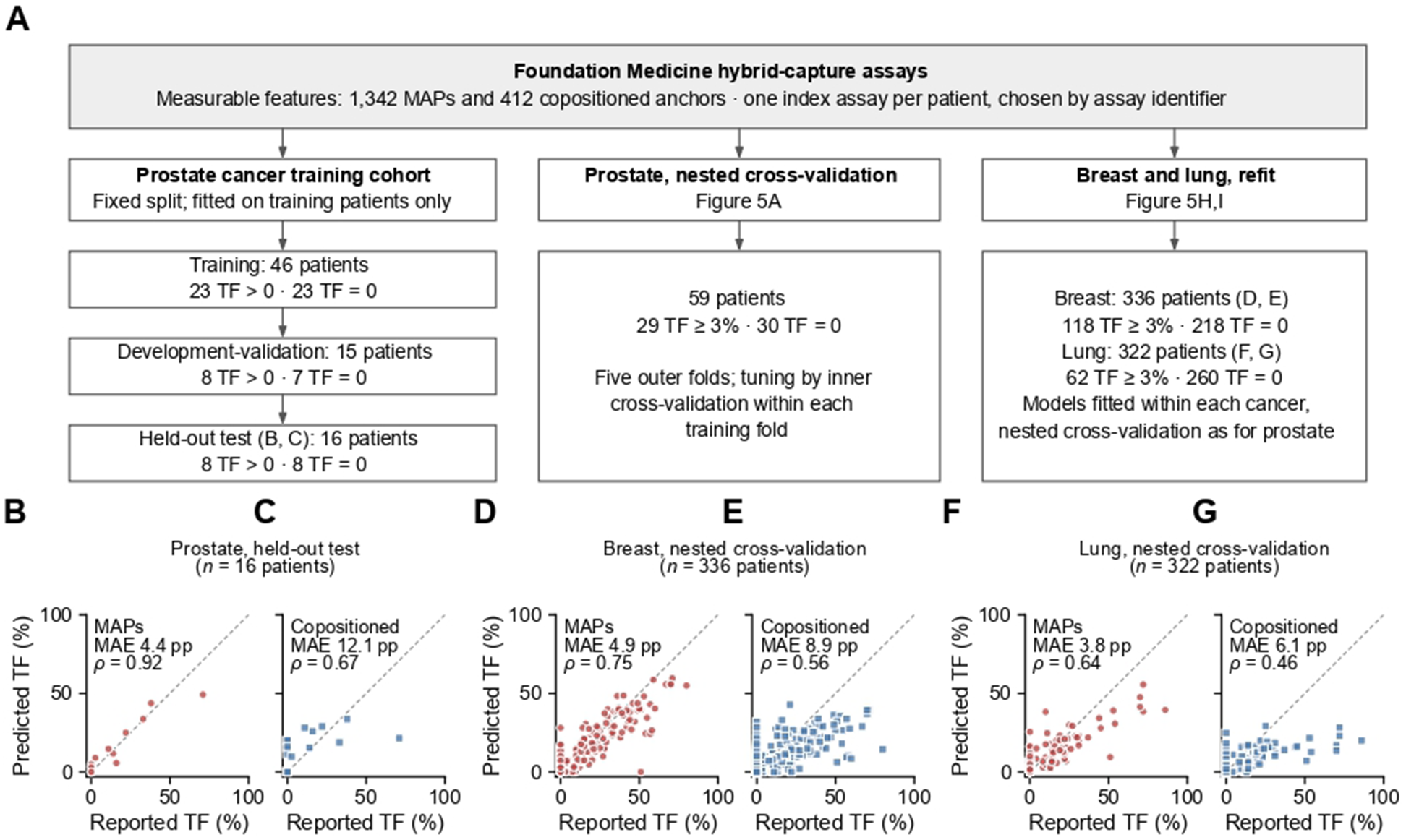
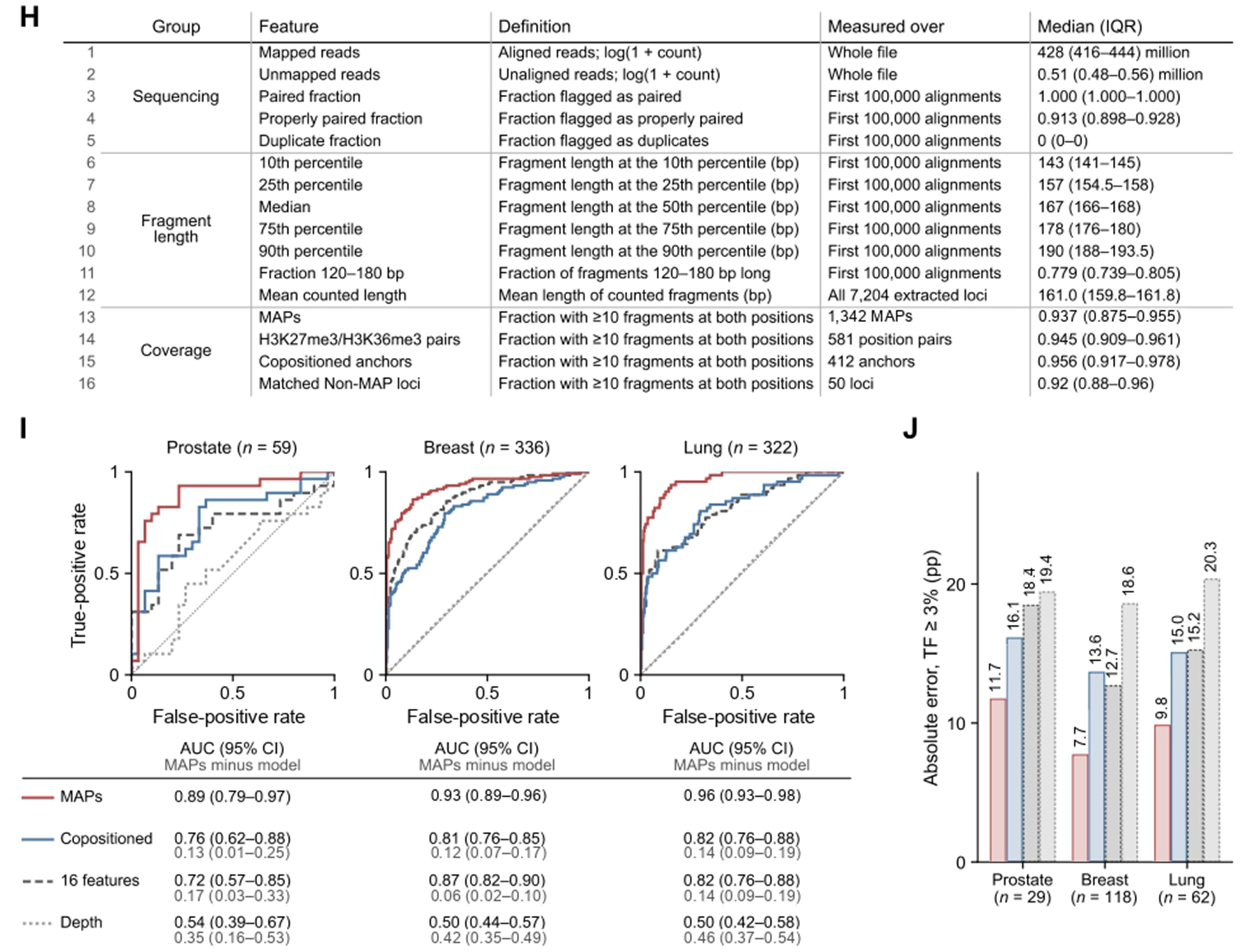
Tumor-fraction estimation in Foundation Medicine specimens: design, prostate test, breast and lung models refit by nested cross-validation, the fragment-length and sequencing features, and the four models compared on the same folds. **(A)** Study design. Paired-position features measurable in Foundation Medicine hybrid-capture specimens comprise 1,342 MAPs and 412 copositioned anchors. Prostate cancer training cohort: one index specimen per patient, chosen by specimen identifier, entered training, development-validation and test groups of 48, 16 and 16 patients; technical eligibility retained 46, 15 and 16, with 23/23, 8/7 and 8/8 reported-positive/reported-zero tumor fraction (TF). Imputation, scaling, cross-validation and fitting used training patients only; the fitted models were applied unchanged to the test patients. Breast and lung models were refit within each cancer by nested cross-validation (**Methods**). **(B, C)** Predicted versus reported TF in the 16 prostate test patients for the MAP **(B)** and copositioned-anchor **(C)** elastic-net models, including all eight reported-zero specimens. **(D–G)** Out-of-fold predicted versus reported TF in breast **(D, E)** and lung **(F, G)** patients with reported TF 0 or ≥3% (336 and 322 patients; 118 and 62 reported-positive), for the MAP and copositioned-anchor models. Each point is one patient; predictions are bounded to 0–100%; dashed lines mark equality; MAE is mean absolute error in percentage points (pp) and ρ the Spearman correlation. **(H)** The 16 fragment-length and sequencing features of the comparator model (I, J; **Figure S11C–E**). Read counts use the whole file; flags and fragment-length percentiles use the first 100,000 primary alignments in coordinate order. Fragment length is the absolute template length of properly paired first reads with same-chromosome mates. Mean counted length uses the 120–180-bp fragments counted at all 7,204 extracted loci (1.23 Mb). A MAP predictor uses only the counted fragments overlapping ±73 bp of its two positions (1,342 MAPs; 0.23 Mb in total). The four coverage features are the fractions of loci in each set with at least ten counted fragments overlapping ±73 bp of both positions. Values are medians (interquartile ranges) among the 59 prostate development patients. **(I)** ROC curves for reported TF 0 versus ≥3% in prostate (59 patients), breast (336) and lung cancer (322) for elastic-net models built on MAPs (red), copositioned anchors (blue), the 16 features of H (gray dashed) and sequencing depth (gray dotted), fitted by nested cross-validation on identical folds; Figure 5C and H show the MAP, copositioned-anchor and depth models. Below, area under the ROC curve (AUC) with 95% patient-bootstrap confidence intervals and, for each comparator, the AUC of MAPs minus that of the comparator (gray, under each AUC) with its 95% paired patient-bootstrap confidence interval. **(J)** Mean absolute error in patients with reported TF ≥3% (prostate, 29; breast, 118; lung, 62) for the same four models, from out-of-fold predictions bounded to 0–100%.

**Figure S10.**
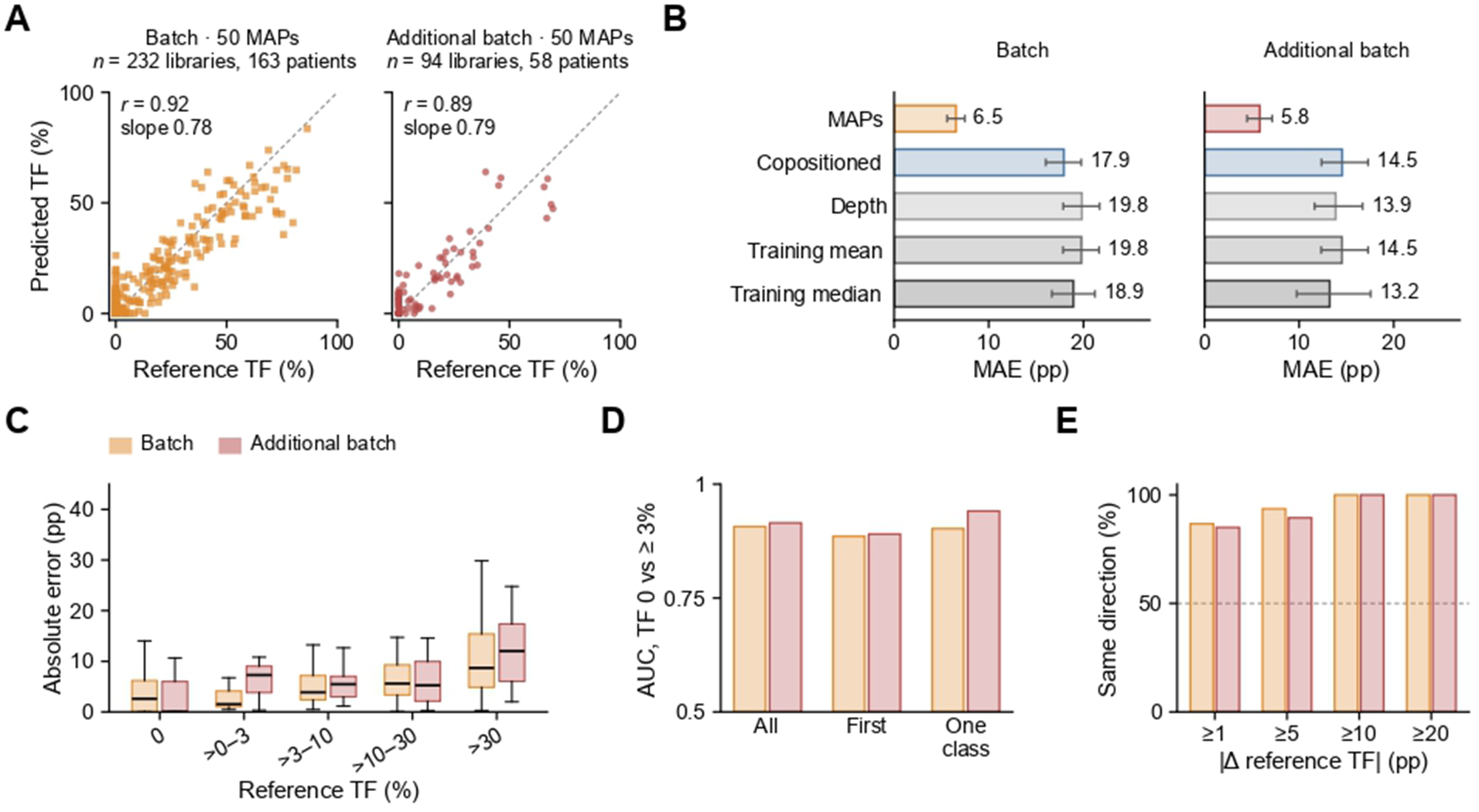
Virtanen–Kwan cohort: two capture batches held out and scored with fixed models. Virtanen–Kwan cohort (Virtanen, Kwan et al., Ann. Oncol., 2026; EGAD50000001594); reference tumor fraction (TF) is the ctDNA percentage in the source study’s Supplementary Table S1. Patients were split into development, tuning and held-out groups before any model was fitted, with every library of a patient in one group. The capture batch and the additional capture batch comprise held-out patients sequenced with the two capture panel generations that cover at least 50 measurable MAPs (≥10 fragments of 120–180 bp at both positions in development libraries). The capture batch uses 50 of its 99 measurable MAPs and the additional batch 50 of its 62, each chosen on tuning patients before held-out scoring. Each elastic-net model was fitted on development patients sequenced with the same panel generation and applied unchanged, with no refitting or recalibration. The batches were scored separately and share no libraries; five patients contributed different blood draws to both batches. **(A)** Predicted versus reference TF for all held-out libraries (capture batch, 232 libraries from 163 patients; additional batch, 94 libraries from 58 patients). r is the patient-weighted Pearson correlation; the slope comes from regressing predicted TF on reference TF. **(B)** Held-out mean absolute error (MAE, in percentage points, pp) with 95% patient-bootstrap CIs for the MAP models and four comparators: a copositioned-anchor model (which gave the same prediction for every library in the additional batch), sequencing depth, and the training-set mean and median. **(C)** Absolute error by reference TF (the group above 0 and below 3% holds three libraries in each batch). Box plots show the median (center line) and the interquartile range (box); whiskers extend to the most extreme values within 1.5 times the interquartile range. **(D)** Area under the receiver operating characteristic curve (AUC) for reference TF 0 versus ≥3%, computed over patients with at least one library at 0 or ≥3%: all libraries (patient-weighted; 160 and 57 patients), the first library per patient in the source’s sample order (160 and 57) and patients whose libraries all fall in one class (152 and 49). **(E)** Paired draws from the same patient, in the source’s sample order: patient-weighted percentage of pairs in which the predicted change has the same sign as the reference change, by minimum reference change of 1, 5, 10 and 20 pp (capture batch, 58, 49, 38 and 20 pairs; additional batch, 20, 19, 15 and 11 pairs).

**Figure S11.**
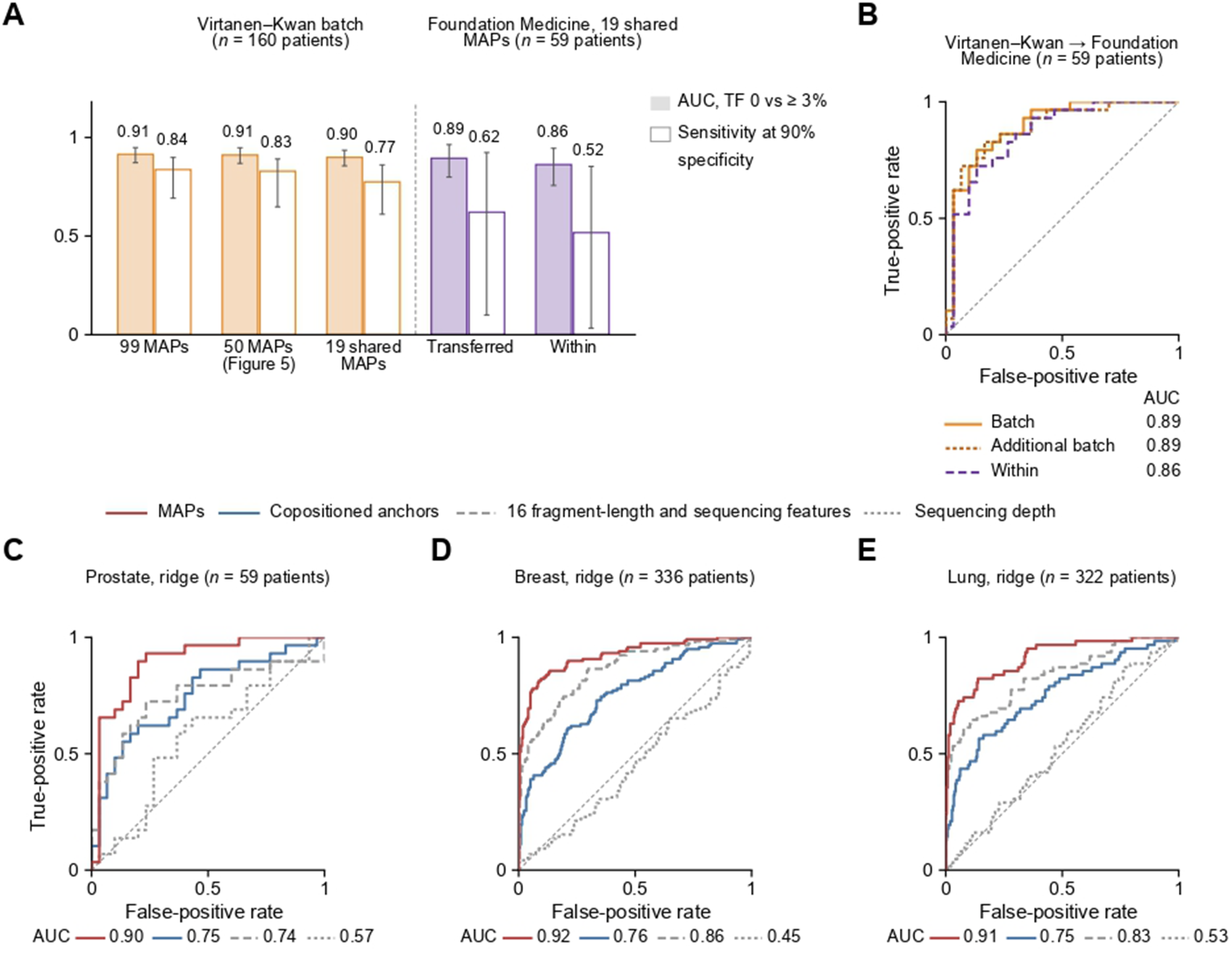
Virtanen–Kwan models with all measurable MAPs and on the MAPs shared with Foundation Medicine, and Foundation Medicine models fitted by ridge regression. **(A)** Area under the receiver operating characteristic curve (AUC; filled) and sensitivity at 90% specificity (open), with 95% patient-bootstrap CIs (2,000 resamples) and each patient weighted equally: the Virtanen–Kwan capture batch scored with the models on all 99 measurable MAPs, on the 50 MAPs of Figure 5D and on the 19 shared MAPs; Foundation Medicine patients scored with the transferred 19-MAP model and with nested cross-validation on the same MAPs. **(B)** ROC curves for the 59 Foundation Medicine prostate patients scored with Virtanen–Kwan models trained on the MAPs measurable on both panels (19 for the capture-batch panel and 18 for the additional-batch panel) and applied unchanged, with imputation and scaling learned in the Virtanen–Kwan cohort, and with Foundation Medicine nested cross-validation on the same 19 MAPs (within). **(C–E)** ROC curves for reported tumor fraction (TF) 0 versus ≥3% in Foundation Medicine patients with prostate **(C)**, breast **(D)** and lung **(E)** cancer, from ridge regressions on MAPs, copositioned anchors, 16 fragment-length and sequencing features and sequencing depth, fitted by nested cross-validation on the same folds as the elastic-net models of Figure 5C and H (out-of-fold predictions). The AUC of MAPs exceeds that of the 16 features by 0.15 (95% patient-bootstrap CI, 0.02–0.29), 0.06 (0.02–0.10) and 0.08 (0.04–0.12).

**Figure S12.**
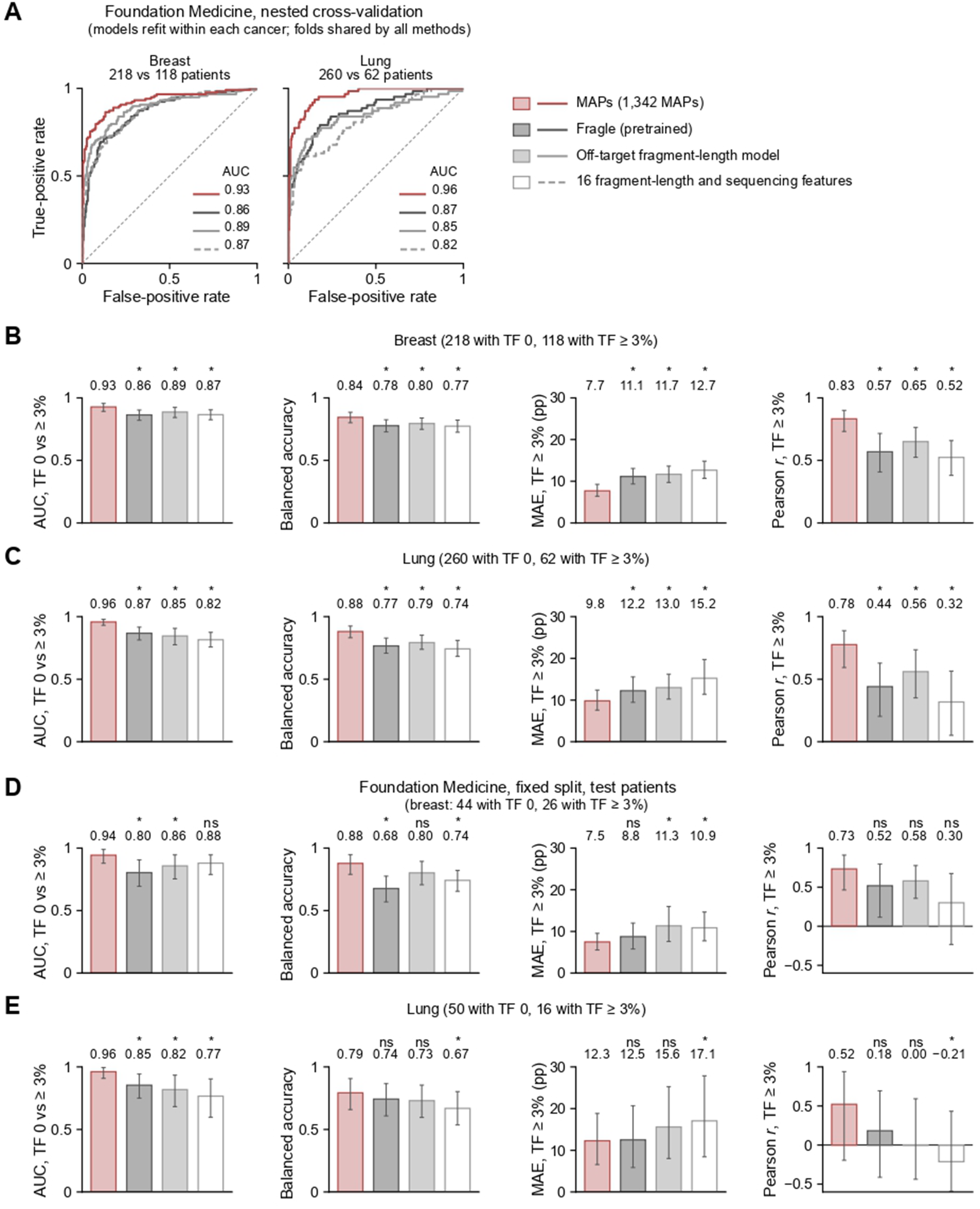
Benchmarking MAPs against Fragle and fragment-length models in Foundation Medicine breast and lung cancer. MAP models refit within each cancer (Figure 5H) compared on the same patients with three fragment-length models: pretrained Fragle, run unchanged in targeted mode on the reads outside an unpadded target mask inferred from coverage; an elastic net on the lengths of the same off-target reads; and the 16 fragment-length and sequencing features of **Figure S9H**. Every threshold and linear calibration came from outer-training (A–C) or validation (D, E) patients (**Methods**). **(A)** ROC curves for reported tumor fraction (TF) 0 versus ≥3% from nested cross-validation with five outer and five inner folds shared by all models (out-of-fold predictions), in breast (336 patients; 218 with TF 0 and 118 with TF ≥3%) and lung cancer (322 patients; 260 and 62); AUCs are listed in each plot. **(B, C)** Nested cross-validation in breast **(B)** and lung cancer **(C)**. **(D, E)** Fixed split: models fitted in training patients, with thresholds and calibration set in validation patients, applied unchanged to test patients with breast (**D**; 70 patients, 44 with TF 0 and 26 with TF ≥3%) or lung cancer (**E**; 66 patients, 50 and 16). Endpoints in B–E: area under the receiver operating characteristic curve (AUC) for TF 0 versus ≥3%; balanced accuracy at the threshold that maximized it in outer-training or validation patients; mean absolute error (MAE, percentage points [pp]) and Pearson *r* in patients with TF ≥3%, from predictions bounded to 0–100%. Bars show estimates with 95% patient-bootstrap confidence intervals (1,000 resamples; each patient weighted equally). The estimate is printed above each bar; above that, * marks a comparator whose paired 95% confidence interval for the MAPs-minus-comparator difference excludes zero, and ns one whose interval includes zero.

**Figure S13.**
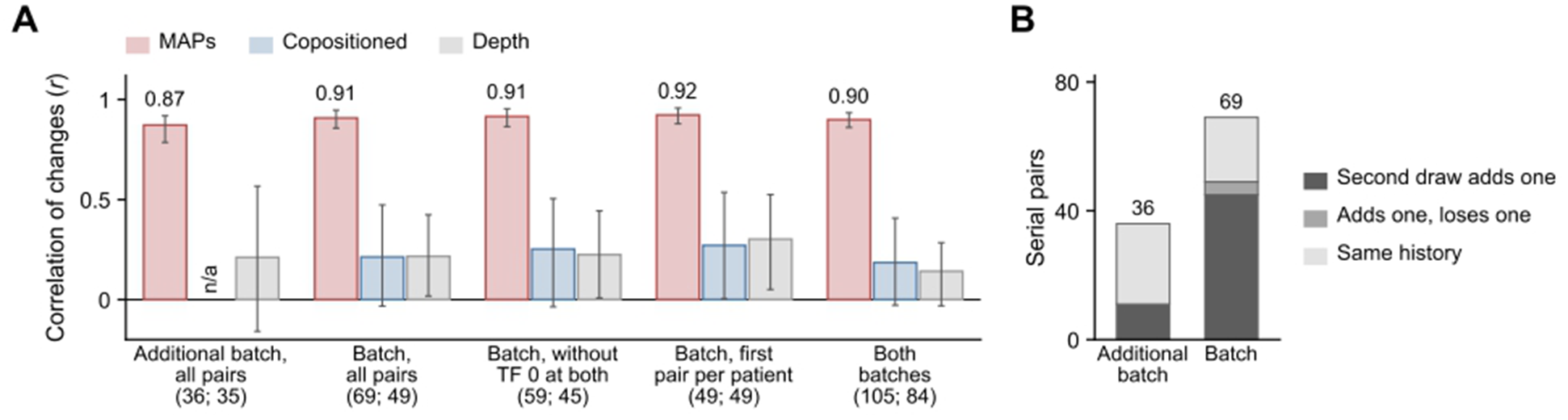
Serial tracking in the Virtanen–Kwan cohort. Virtanen–Kwan cohort; reference tumor fraction (TF) is the ctDNA percentage in the source study’s Supplementary Table S1. The capture batch and the additional capture batch are the held-out batches of **Figure S10** (the capture batch is the held-out set of Figures 5 **and 6**), scored with fixed models. **(A)** Agreement between the change in predicted and the change in reference TF across paired draws from the same patient (taken in the source’s sample order): patient-weighted Pearson r with 95% patient-bootstrap CIs (1,000 resamples) for MAPs, copositioned anchors and sequencing depth, in the additional batch, in the capture batch (all pairs; excluding pairs with reference TF 0 at both draws; the first pair per patient) and in both batches together. Displayed pair and patient counts apply to the MAP and depth models. The copositioned model had 66, 56 and 48 evaluable pairs in the three capture-batch analyses, respectively, and 102 across batches. In the additional batch the copositioned model gave the same prediction for every library, so its r is undefined (n/a). **(B)** Treatment history between the two draws of each pair, from the source table: the second draw adds a treatment (additional batch, 11 pairs; capture batch, 45), adds one and loses another (0; 4) or has the same history (25; 20). No pair only loses a treatment, consistent with the sample order.

---

**Supplementary Note Table 1a.** Within-cohort pairing. Additional pairs (%) = 100 × (nearest-partner assignments − reciprocal pairs)/reciprocal pairs.

| Window (bp) | Cohort | Nearest-partner assignments | Reciprocal pairs | Additional pairs (%) |
| --- | --- | --- | --- | --- |
| 50 | 1 | 897,669 | 897,569 | 0.01 |
| 50 | 2 | 1,493,595 | 1,493,046 | 0.04 |
| 73 | 1 | 1,011,192 | 1,006,564 | 0.46 |
| 73 | 2 | 1,983,906 | 1,959,804 | 1.23 |
| 100 | 1 | 1,118,191 | 1,095,900 | 2.03 |
| 100 | 2 | 2,442,731 | 2,342,358 | 4.29 |
| 150 | 1 | 1,353,355 | 1,242,877 | 8.89 |
| 150 | 2 | 3,079,472 | 2,768,778 | 11.22 |

**Supplementary Note Table 1b.** Cross-cohort matching at a 73-bp within-cohort pairing window. Direction-eligible pairs have at least 10-bp absolute separation in both cohorts; agreement is the percentage with the same upstream–downstream order.

| Tolerance (bp) | Matched pairs | Direction-eligible pairs | Same order | Opposite order | Agreement (%) |
| --- | --- | --- | --- | --- | --- |
| 20 | 91,471 | 38,627 | 26,810 | 11,817 | 69.4 |
| 30 | 120,489 | 53,131 | 35,120 | 18,011 | 66.1 |
| 50 | 161,975 | 76,137 | 45,605 | 30,532 | 59.9 |

## METHODS

### Participants and cohorts

#### Human participants and ethics

All research involving human participants, specimens or data was conducted in accordance with the Declaration of Helsinki. Plasma collection and profiling for the NCI discovery cohorts were approved by the Institutional Review Board of the National Institutes of Health as part of a multi-cohort phase 2 trial (ClinicalTrials.gov Identifier: NCT02484404, “Phase I/II Study of the Anti-Programmed Death Ligand-1 Durvalumab Antibody (MEDI4736) in Combination With Olaparib and/or Cediranib for Advanced Solid Tumors and Advanced or Recurrent Ovarian, Triple Negative Breast, Lung, Prostate and Colorectal Cancers”, PI: Jung-Min Lee), first registered on June 25, 2015.^68^ This study was conducted in accordance with ethical principles that have their origin in the Declaration of Helsinki and are consistent with the International Council on Harmonization guidelines on Good Clinical Practice, all applicable laws and regulatory elements, and all conditions required by a regulatory authority and/or institutional review board. Written informed consent was obtained from all patients prior to performing study-related procedures in accordance with federal and institutional guidelines. Specimens of the independent cohort and its inter-laboratory replicates were collected under University of British Columbia Research Ethics Board certificates H18-00944 and H16-00934 with written informed consent.^21^ Specimens of the Baca cohort were collected under Dana-Farber/Harvard Cancer Center protocol 01-045 and National Cancer Institute protocols 08-C-0074 and 10-C-0062.^20^ Specimens of the DFCI NEPC-monitoring cohort were identified from the Dana-Farber Cancer Institute Arthur and Linda Gelb Center for Translational Research biobank and clinical database under Dana-Farber/Harvard Cancer Center protocol 01-045 and were de-identified before analysis. Specimens of the Emory capture cohort were collected under institutional review board protocol 2025P010474. Published and commercial datasets were analyzed under the access terms of their sources (Data availability).

#### Discovery cfChIP-seq cohort

The discovery cohorts comprised 37 serial plasma specimens from 15 participants with metastatic castration-resistant prostate cancer that had progressed on androgen receptor-targeted therapy, enrolled in a clinical trial of durvalumab and olaparib (NCT02484404)^68^ and profiled in the laboratory of D. Y. Takeda (National Cancer Institute). Discovery cohort 1 comprised 17 specimens from seven participants and discovery cohort 2 comprised 20 specimens from eight participants, collected at baseline, restaging and progression. No participant overlapped with the cohort of Sipola et al.^21^ Thirty-six specimens had both H3K4me2 and H3K9me3 profiles; one specimen of discovery cohort 2 had an H3K4me2 profile only and was excluded from paired-mark analyses. Tumor fraction was estimated with ichorCNA^42^ from the unenriched input library of each specimen.

#### Independent cfChIP-seq cohort and inter-laboratory replicates

The independent cohort comprised the ten plasma specimens of Sipola et al.^21^ with both H3K4me2 and H3K9me3 profiles (patients P3, P6, P8, P12, P13, P19, P24, P29, P30 and P34; Supplementary Table 2 of that study): nine from patients with metastatic prostate cancer (ctDNA fraction 49.6–84.6%) and one from a patient with metastatic urothelial carcinoma. Participants were enrolled in a province-wide plasma biobanking program at the Vancouver Prostate Centre and BC Cancer (British Columbia, Canada), with plasma collected before a new line of systemic therapy and at progression. Of the patients with prostate cancer in that study, 85% had metastatic castration-resistant disease at first collection, 47% had received at least one androgen receptor-targeted therapy and 17% at least one taxane.^21^

Inter-laboratory replicates were new libraries from seven of the ten specimens (P3-1, P6-2, P12-2, P19-1, P24-1, P29-1 and P34-1), prepared independently by the Takeda and Wyatt laboratories with the protocol of Sipola et al.^21^ (Plasma cfChIP-seq) for H3K4me2, H3K9me3, H3K27me3 and H3K36me3, each with an unenriched input library. Neither the independent cohort nor its inter-laboratory replicates were used for MAP discovery.

#### Deep plasma cfDNA WGS cohort

Deep plasma cfDNA WGS of Herberts et al.^44^ comprised 48 specimens from 30 patients with metastatic castration-resistant prostate cancer, sampled before and after progression on androgen receptor pathway inhibitors, with tumor fraction taken from Supplementary Table 1 of that study (’WGS cancer fraction’). Six control libraries were assigned a tumor fraction of zero: four from Snyder et al.^1^ (BH01, from pooled healthy plasma, and IH01, IH02 and IH03) and two designated as controls by Herberts et al.^44^ (CTRL-AE-179 and CTRL-AE-201).

#### Emory cfDNA capture cohort

The Emory capture cohort comprised 33 unpublished plasma cfDNA specimens sequenced by hybrid capture in the laboratory of J. E. Berchuck (Winship Cancer Institute of Emory University): 26 specimens from 16 patients with metastatic castration-resistant prostate cancer (mCRPC) and seven specimens from seven cancer-free controls. cfDNA was subjected to Illumina paired-end sequencing after hybrid capture with custom baits generated by Twist Biosciences for a separate project; MAPs were not targeted by the panel. After filtering (Hybrid-capture sequencing), specimens had a median of 35.8 million fragments (range 16.5–59.3 million). Figure 5A and Figure S8 used all 33 specimens; Figure 5B used an earlier release of 22 specimens from 13 of the patients with mCRPC and the seven controls.

#### Virtanen–Kwan cfDNA capture cohort

The Virtanen–Kwan cohort comprised 2,700 targeted plasma cfDNA libraries, with matched leukocyte DNA, from 1,496 patients with metastatic castration-resistant prostate cancer (Virtanen, Kwan et al.;^55^ European Genome-phenome Archive, EGAD50000001594). Participants were enrolled through prospective clinical trials and the Genitourinary Cancer Biobank of BC Cancer (British Columbia, Canada), and specimens were sequenced with four successive generations of a targeted panel. Reference tumor fraction (ctDNA percentage) and prior treatment at each draw were taken from Supplementary Table S1 of that study. Before any model was fitted, patients were assigned to development (n = 898), tuning (n = 299) and held-out (n = 299) sets, with all libraries of a patient in one set.

Of the 232 held-out libraries used in Figures 5 and 6, 78% came from patients with prior androgen-deprivation therapy, 39% with a prior androgen receptor pathway inhibitor and 17% with a prior taxane; entries recorded as ‘Partial’ were not counted. Sample names carry a sequential draw number (for example, 0005-cfDNA-5th) but no collection date, and serial libraries of a patient were ordered by this number.

#### Foundation Medicine cfDNA capture cohort

De-identified Foundation Medicine cfDNA data, all from one version of the FoundationOne Liquid CDx panel (bait set AB1) and each with a reported tumor fraction, were obtained for patients with prostate adenocarcinoma treated at Dana-Farber Cancer Institute (89 eligible specimens from 80 patients), breast cancer (389 patients) and lung cancer (378 patients). Of the 767 patients with breast or lung cancer, 614 had a recorded histology: all 311 breast cancers were invasive breast carcinoma without further subtype, and the 303 lung cancers were adenocarcinoma (167), non-small-cell lung carcinoma not otherwise specified (32), lung carcinoma not otherwise specified (87), squamous cell carcinoma (8), small-cell carcinoma (5) or other histologies (4). Specimens with a reported tumor fraction of 0% came from patients with cancer and formed the negative class (0% versus ≥3%). Treatment information was not available.

#### Prostate cancer low-pass cfDNA WGS cohorts

The Baca cohort^20^ comprised published low-pass plasma cfDNA WGS of 32 patients with prostate cancer, one specimen per patient: 24 with prostate adenocarcinoma (PRAD) and eight with neuroendocrine prostate cancer (NEPC), as annotated in that study.

The DFCI NEPC-monitoring cohort comprised 39 low-pass plasma WGS specimens from nine participants with prostate cancer followed for neuroendocrine transformation (Gelb Center biobank; G. G. Lee et al., manuscript in preparation). One further participant was excluded because a specimen from this participant had been profiled in a prostate cancer hybrid-capture dataset examined during method development. The cohort investigators annotated each specimen as adenocarcinoma, pre-NEPC or NEPC. Specimens annotated adenocarcinoma or pre-NEPC and collected before a participant’s first NEPC-labeled specimen formed the earlier state (16 specimens) and were compared with the 23 NEPC-labeled specimens. Six participants had specimens in both states.

### Laboratory methods

#### Plasma cfChIP-seq

Plasma collection, cfDNA isolation and nucleosome immunoprecipitation for the discovery cohorts, and nucleosome immunoprecipitation for the inter-laboratory replicates, followed Sipola et al.^21^ Blood was collected in Streck Cell-Free DNA BCT tubes, and plasma was separated according to the manufacturer’s instructions and stored at −80 °C. Cell-free nucleosomes were immunoprecipitated directly from plasma. Antibodies against H3K4me2 (Diagenode, C15410035), H3K9me3 (Diagenode, C15410193), H3K27me3 (Diagenode, C15410195) and H3K36me3 (Diagenode, C15410192) were coupled to Dynabeads M-270 Epoxy (Thermo Fisher Scientific, 14302D) at 5 µg of antibody per mg of beads, and coupling was confirmed by immunoblotting of the supernatant before and after coupling. Antibody-coupled beads were added to 300 µl–2 ml of plasma, depending on cfDNA concentration, and incubated overnight at 4 °C; a fraction of the same plasma was kept before immunoprecipitation as unenriched input. Beads were washed three times with buffer A (50 mM Tris pH 7.5, 2 mM EDTA, 150 mM NaCl, 0.1% sodium deoxycholate, 1% Triton X-100), three times with buffer B (50 mM HEPES pH 7.6, 1 mM EDTA, 500 mM LiCl, 0.7% sodium deoxycholate, 1% NP-40) and twice with TE (10 mM Tris pH 8.0, 1 mM EDTA). DNA was purified after proteinase K and RNase treatment with the Monarch DNA Cleanup kit (New England Biolabs). Libraries from immunoprecipitated and input DNA were prepared with the NEBNext Ultra II Library Preparation kit (New England Biolabs) and sequenced on an Illumina NextSeq 2000 at the CCR Genomics Core, National Cancer Institute.

#### MNase–ChIP-seq of LNCaP cells

LNCaP cells (ATCC; authenticated by short tandem repeat profiling and free of mycoplasma) were washed in ice-cold PBS with protease inhibitors, scraped and pelleted. Nuclei from 5 × 10⁶ cells were released in 200 µl of digestion buffer (50 mM Tris-HCl pH 7.6, 1 mM CaCl₂, 0.2% NP-40, 5 mM sodium butyrate and protease inhibitors) with a Dounce homogenizer and digested at 37 °C with 0.6 µl of micrococcal nuclease (0.04 U/µl; Sigma-Aldrich, N3755), and aliquots were stopped with EDTA-containing buffer at 2, 4, 6 and 8 min. Digests were briefly sonicated, diluted in ChIP lysis buffer, checked for mononucleosomes (about 146 bp) on an Agilent 4150 TapeStation and cleared by centrifugation. Nucleosomes were immunoprecipitated as described,^20^ with antibodies against H3K4me2 (Diagenode, C15410035) or H3K9me3 (Diagenode, C15410193) bound to Dynabeads Protein A/G (Invitrogen), and washed in low-salt, high-salt and LiCl buffers. DNA was released with proteinase K and RNase A and purified with the ChIP DNA Clean & Concentrator kit (Zymo Research, D5205). Libraries were prepared with the ThruPLEX DNA-Seq kit (Takara Bio, R400675) and sequenced (150-bp paired-end) on an Illumina NovaSeq 6000 at Novogene.

### Identification of MAPs

#### Sequencing data processing

Libraries of the discovery and independent cohorts were processed with SNAPIE (v1.5.1),^69^ a Nextflow-based^70^ pipeline for cfChIP-seq, and the inter-laboratory replicates with SNAPIE v1.5.18, which uses the same alignment and filtering settings. Adapters were trimmed with Trim Galore (v0.6.10 and v0.6.7, respectively; https://github.com/FelixKrueger/TrimGalore) and Cutadapt^71^ (v3.4 for the inter-laboratory replicates) with quality trimming disabled (--quality 0) to preserve fragment ends. Reads were aligned to hg19 (GCA_000001405.1) with BWA-MEM (v0.7.18)^72^ using a fixed insert-size distribution (mean 250 bp, standard deviation 200 bp, maximum 2,000 bp and minimum 10 bp). Alignments were sorted with SAMtools (v1.20.1),^73^ and duplicates were removed with Picard MarkDuplicates (v2.27.4; https://broadinstitute.github.io/picard/). Properly paired reads with mapping quality ≥30 were retained, and fragments were extracted with BEDTools (v2.30.0).^74^ Fragments overlapping the ENCODE hg19 exclusion list (ENCFF001TDO),^75^ shorter than 20 bp or longer than 500 bp were removed. Library quality was summarized by retained fragment yield, fragment-length distribution (Figure S2A) and mononucleosomal fragment frequency; no library was excluded on these summaries.

#### Genome builds

MAP and copositioned-anchor coordinates were defined and are reported in hg19 (GRCh37). Three datasets were analyzed in hg38 (GRCh38): the deep WGS of Herberts et al.^44^ and the Virtanen–Kwan cohort,^55^ which were released aligned to hg38, and the Emory capture cohort, which was aligned to hg38 (Hybrid-capture sequencing). For these datasets, both positions of each MAP were converted separately from hg19 to hg38 with pyliftover (v0.4.1; https://github.com/konstantint/pyliftover) and the UCSC hg19ToHg38 chain file,^76^ keeping the first mapping, and the pairing, class and orientation of each MAP were kept from discovery; all 8,044 MAPs of the deep WGS score converted. All other datasets were analyzed in hg19.

#### Nucleosome dyad calling

Nucleosome dyads were called separately for H3K4me2 and H3K9me3 from fragments pooled across the specimens of each discovery cohort and were retained only when two callers agreed. Probabilistic nucleosome scores (PNS) were calculated with the implementation of Johnston et al.^45^ (https://github.com/traulab/PNS; commit b182807) using a nucleosome length of 167 bp, the modal length of plasma mononucleosomal fragments, and fragment-length bounds of 127–207 bp, and the center of each local PNS peak was taken as a candidate dyad. Candidates with normalized PNS ≥5 were retained when DANPOS3 dpos^46^ (https://github.com/sklasfeld/DANPOS3; commit cb10355; default parameters, paired-end mode, -m 1), run separately for each mark, called a dyad within 20 bp. The PNS peak center was used as the dyad coordinate (Figure S1A–C).

#### Definition of MAPs and copositioned anchors

H3K4me2 and H3K9me3 dyads were paired within each cohort when they were reciprocal nearest neighbors no more than 73 bp apart, about half of the 147-bp nucleosome core,^47,48^ within which two dyads share more than half of their core DNA (Figure S1D,F; Supplementary Note 1).

Pairs were matched between cohorts when their H3K4me2 dyads were reciprocal nearest neighbors within 30 bp (Figure S1E). The H3K4me2-associated position of each locus was the integer midpoint of the H3K4me2 dyads of the two cohorts, and the H3K9me3-associated position the integer midpoint of their H3K9me3 dyads.

MAPs were loci with the same order of the two marks and a separation of 10–73 bp in both cohorts (35,120 MAPs). The 10-bp lower bound is about one helical turn of DNA (10.4 bp), at which the two positions keep the same rotational setting.^29,47^ Copositioned K4/K9 anchors were loci separated by 0–4 bp in both cohorts (4,774 loci), less than half a helical turn. Loci separated by 5–9 bp and the other matched loci that met neither definition (80,595 loci) were termed Non-MAP loci and used as matched comparators (Figure 3D; Figure S6F; Figure S9H).

#### Robustness of the MAP definition

##### Copositioned cutoff

The cutoff was varied from 2 to 8 bp, and the percentage of anchors within 1 kb of a transcription start site (TSS; hg19 knownGene) was compared with that of MAPs, with 95% intervals from 5,000 bootstrap samples of chromosomes (Figure S1G–I).

##### Within-cohort pairing distance

Reciprocal nearest-neighbor pairs, and the percentage of pairs more than 73 bp apart, were counted as the maximum pairing distance increased (Figure S1D; Supplementary Note 1).

##### Cross-cohort matching tolerance

Matching was repeated with tolerances of 20 and 50 bp (26,810 and 45,605 MAPs), and the separations of matched loci were compared between cohorts (Figure S1E; Supplementary Note 1).

##### Read depth

PNS profiles were rebuilt from 50%, 75% and 100% of fragments (three random seeds each) at 460 loci chosen without reference to signal (ten MAPs and ten copositioned anchors per chromosome). A peak was recovered when it lay within 20 bp of its full-depth position (Figure S1J).

### Characterization of MAPs

#### Windowed protection score

Protection at each position was measured with the long-fragment windowed protection score (WPS) of Snyder et al.,^1^ computed from fragments of 120–180 bp (fragments of all lengths in the Emory capture cohort). For a 120-bp window centered on coordinate *x*, where *N*_span_ is the number of fragments spanning the window and *N*_end_ the number of fragment ends within it. The MAP contrast was the WPS at the H3K4me2-associated position minus that at the H3K9me3-associated position. Deep WGS used a per-base form of this score, and the normalization to local coverage for each data type is given in its section.

#### Genomic annotation

Each locus was represented by its hg19 midpoint. Gene features and distance to the nearest TSS were assigned with ChIPseeker (v1.42.1)^77^ and TxDb.Hsapiens.UCSC.hg19.knownGene (v3.2.2). Midpoints were intersected with the EpiMap 18-state ChromHMM segmentation of normal prostate gland (BSS01456)^13^ and the SCREEN v4 registry of candidate cis-regulatory elements (cCREs),^78^ including promoter-like (PLS), proximal enhancer-like (pELS, within 2 kb of a TSS^79^), distal enhancer-like (dELS), accessible (CA, CA-H3K4me3, CA-CTCF and CA-TF) and transcription-factor-bound (TF) elements.

Loci more than 1 kb from the nearest TSS were classified as gene body when the midpoint lay within a protein-coding RefSeq gene (UCSC refGene, NM_ transcripts) more than 1 kb from any coding TSS, and as intergenic otherwise (Figure 3A). Enrichment was tested by two-sided Fisher’s exact test, with log₂ odds ratios and chromosome-bootstrap 95% confidence intervals.

For Figure S5A–D, MAPs were divided into six separation strata (10–19 to 60–73 bp), each compared with all copositioned anchors (log₂ odds ratios with a 0.5-cell correction); binary annotations used 151-bp windows centered on the locus midpoint, and chromatin states were assigned at the midpoint.

#### Comparison with copositioned anchors

Copositioned anchors had higher discovery read depth than MAPs, and the two classes were therefore compared within depth strata. The log₂ read depth of each mark, summed over the discovery cohorts, was divided into quartiles across the 39,894 MAPs and copositioned anchors, and the two sets of quartiles were crossed into 16 strata. For each of 59 reference region sets fixed before testing (Supplementary Table 2), the Mantel–Haenszel odds ratio was tested by Cochran–Mantel–Haenszel χ² test with continuity correction and Benjamini–Hochberg adjustment (Figure 3B,C). A host gene was the protein-coding RefSeq gene containing the locus midpoint more than 1 kb from any coding TSS; it was considered expressed in a tissue at a GTEx v8^80^ median TPM ≥1 and broadly expressed when expressed in at least 40 of 54 tissues.

For Figure 3D, a promoter was active across lineages when the locus midpoint lay in a TSS-associated chromatin state in prostate (EpiMap BSS01456: TssA, TssFlnk, TssFlnkU, TssFlnkD or TssBiv), neutrophils (Roadmap E030) and monocytes (Roadmap E029; 1_TssA, 2_TssAFlnk, 10_TssBiv or 11_BivFlnk).^17^ A locus was CTCF-bound in prostate and blood when an LNCaP CTCF peak and a K562, GM12878 or neutrophil CTCF peak lay within 150 bp of either position. Both features entered one logistic regression of MAPs versus copositioned anchors together with chromatin activity restricted to one or two of the three cell types, adjusted for z-scored log₂ H3K4me2 and H3K9me3 depth, GC fraction and mappability. Standard errors were clustered on 5-Mb genomic blocks, and P values came from Wald tests.

#### Binding sites and dyad separation

A locus was near a binding site when a ChIP-seq peak lay within 150 bp of either position. Peak sets were CTCF in LNCaP (ENCODE ENCSR315NAC; conservative IDR peaks ENCFF685MPH), prostate tissue (present in at least five of ten experiments) and neutrophils; RAD21 in GM12878 and K562; AR, FOXA1 and H3K27ac in localized prostate tumors and normal prostate from Pomerantz et al.;^81^ AR from recurrent tumor sites;^82^ HOXB13 in prostate tumors; and LNCaP ABC candidate elements.^53^ LNCaP CTCF peaks were divided into thirds by IDR signal value, each locus taking the strongest peak within 150 bp, and blood CTCF peaks were the union of K562 (ENCSR000EGM) and GM12878 (ENCSR000AKB) peaks (Figure 3C).

Within MAPs, logistic regression tested whether the odds of a nearby peak changed with separation, measured as the mean absolute offset across the discovery cohorts (odds ratio per 10 bp). The test was repeated under eight prespecified conditions: (1) all loci; (2) loci more than 1 kb from a TSS; (3) loci more than 1 kb from a TSS and outside CpG islands; (4) all loci adjusted for promoter proximity, CpG-island overlap, GC content, mappability and discovery-coverage quintile; (5, 6) separation from discovery cohort 1 or 2 alone; and (7, 8) odd or even chromosomes. A factor decreased with separation when the odds ratio was below one in all eight conditions, the Holm-adjusted P value was below 0.05 in conditions 1 and 4, and P was below 0.05 in conditions 5 and 6 (Supplementary Table 3).

#### Enhancer–gene links

MAP and copositioned coordinates from discovery cohort 1 (cohort 2 in a repeat analysis) were matched to LNCaP candidate elements of the Activity-by-Contact (ABC) model,^53^ one element per eligible locus. Among elements not classed as promoters, a locus had a predicted enhancer–gene link when its element took part in at least one distal enhancer–gene prediction with an ABC score of at least 0.022 (Figure 3E; Figure S5E).

#### Tumor fraction-correlated MAPs

H3K4me2 and H3K9me3 cfChIP-seq signals at each of the 35,120 MAPs, scaled to the median library, were correlated with ichorCNA tumor fraction across the 36 discovery specimens (Spearman ρ). K4/TF-correlated MAPs had H3K4me2 ρ > 0.2 and H3K9me3 ρ < −0.2 (1,776 loci), K9/TF-correlated MAPs the converse (1,619 loci), and tumor fraction-neutral MAPs |ρ| < 0.1 for both marks (3,925 loci). To set the cutoff, each MAP was paired within chromosome with a Non-MAP locus matched on discovery coverage of both marks in both cohorts, GC content and mappability, and the percentage of loci whose two marks correlated with tumor fraction in opposite directions beyond 0.1, 0.2 or 0.3 was compared (exact McNemar test; Figure S6F). At 0.2, MAPs exceeded matched loci and each group kept more than 1,000 MAPs.

Host genes were protein-coding genes containing a MAP in the gene body, excluding genes with both K4/TF-correlated and K9/TF-correlated MAPs. Tumor-upregulated genes had median expression in 497 TCGA primary prostate tumors^83^ at least 1 log₂ unit (log₂(RSEM + 1)) above that in 52 normal prostate samples. Blood-enriched genes had a GTEx v7^84^ whole-blood median TPM ≥1 and at least four times the mean of the median TPMs of all other tissues. Each group was compared with the other MAPs for these gene classes and for proximity to AR and FOXA1 peaks in prostate tumors by two-sided Fisher’s exact test (Figure 4B).

#### Validation in independent plasma

Discovery coordinates, classes and orientations were held fixed in the independent cohort and its inter-laboratory replicates. For each mark, midpoints of 120–180-bp fragments within ±110 bp of the H3K4me2-associated position were summed over three panels of 4,000 loci, drawn in a fixed pseudo-random order from the 4,774 copositioned anchors, the 27,798 MAPs whose separations, averaged over the two discovery cohorts, were 10–40 bp (median 25 bp) and the 7,322 MAPs more than 40 bp apart (40.5–72.5 bp; median 47 bp). Profiles were averaged over loci, divided by the number of 120–180-bp fragments of the library in millions, oriented from the H3K4me2-toward the H3K9me3-associated position and smoothed with a 9-bp running mean.

In each specimen and panel, the separation between the two marks was the shift of the H3K9me3 profile against the H3K4me2 profile, in 1-bp steps from −73 to +73 bp over ±90 bp of standardized profiles, that gave the highest Pearson correlation (Figure 2C). Each MAP group was compared with copositioned anchors within specimen by exact one-sided paired sign test with Holm correction across the four group-by-cohort comparisons, and the two MAP groups were compared within each cohort by exact two-sided sign test.

For individual loci, dyads were called in each validation cohort with the discovery rules (PNS ≥5 and a DANPOS3 call within 20 bp) and paired when the H3K9me3 dyad lay within 73 bp of the H3K4me2 dyad. Each discovery locus was matched to the nearest external pair whose H3K4me2 dyad lay within 30 bp of its H3K4me2-associated position, and the external separation (H3K9me3 minus H3K4me2 dyad position) was compared between the independent cohort and its inter-laboratory replicates by two-sided Mann–Whitney test (Figure 2D). Among matched MAPs with an external separation of at least 10 bp, the percentage with the discovery order of the two marks was tested against 50% by one-sided exact binomial test (Figure 2E).

#### Additional histone marks

H3K27me3 and H3K36me3 cfChIP-seq of the inter-laboratory replicates was read at the fixed positions. In each specimen, the MAP contrast was summed over MAPs with fragments at both positions, divided by the summed number of fragments overlapping the two 120-bp windows (averaged over the two positions) and expressed per 100 fragments. The same value computed in the input library from the same plasma was subtracted, and input-corrected H3K36me3 and H3K27me3 contrasts were compared within specimen by exact two-sided sign test (Figure 2F,G).

Side preference (Figure S4) was (R − L)/(R + L), where R and L are the numbers of fragment centers 10–73 bp from the H3K4me2-associated position on the H3K9me3 side and on the opposite side, counted over all MAPs or all copositioned anchors. For H3K27me3 and H3K36me3, the value of the same-specimen input was subtracted, and MAPs and copositioned anchors were compared by exact two-sided Wilcoxon signed-rank test; for H3K4me2 and H3K9me3, the summed input of the inter-laboratory replicates was subtracted from the merged independent-cohort libraries.

#### MNase–ChIP-seq analysis

Native MNase–ChIP-seq of vehicle-treated LNCaP cells was obtained from Chen et al.^50^ (GSE43720; H3K4me2, GSM1069675 and GSM1069676; H3K9me3, GSM1069679 and GSM1069680; MNase digestion for 10–20 min; 35-bp single-end reads aligned to hg19). Fragment centers were placed 73 bp from the 5′ end of each read on its strand, assuming a fragment length of 147 bp; because reads on the two strands are shifted in opposite directions, an error in this length broadens the averaged profile without moving its center. For each replicate, centers were counted from −100 to +100 bp of the H3K4me2-associated position at MAPs and copositioned anchors, oriented toward the H3K9me3-associated position and expressed as a percentage of all centers in the window, and the two replicates of each mark were averaged (Figure 2H).

MNase–ChIP-seq generated in this study was analyzed in the same way from the centers of 120–180-bp paired-end fragments (hg19; Figure 2I). The percentage of centers on the H3K9me3 side was R/(R + L) × 100, with R and L as defined above.

### Tumor fraction and cancer state from cfDNA

#### Deep plasma WGS: protection at MAP positions

Deep WGS of Herberts et al.^44^ was analyzed from the released hg38 alignments (Bowtie 2) at MAP positions converted to hg38 (Genome builds; all 8,044 MAPs). For each MAP, a per-base protection profile over 501 bp centered on the H3K4me2-associated position was built from 120–180-bp fragments of properly paired, non-duplicate primary alignments with mapping quality ≥30: each fragment added one at every base it covered and subtracted one at each of its two end bases. Profiles with a positive mean were divided by their mean and smoothed with a Gaussian kernel (σ = 5 bp). Protection at a position was the mean of the profile within 15 bp of it, and the MAP contrast was the protection at the H3K4me2-associated position minus that at the H3K9me3-associated position.

#### Deep plasma WGS: MAP score

MAPs and their orientations were fixed from discovery cfChIP-seq before any WGS library was scored. A MAP was selected when its H3K4me2 and H3K9me3 signals, with each library’s counts scaled to the median library total over the 35,120 MAPs, correlated with ichorCNA tumor fraction in opposite directions across the 36 discovery specimens (|ρ| > 0.1 for both marks; 8,257 loci), and was kept when the opposite signs held after counts were normalized over all 120,489 matched loci (8,044 MAPs: 4,222 at which H3K4me2 and 3,822 at which H3K9me3 rose with tumor fraction). Each contrast was oriented toward the position whose mark rose with tumor fraction.

The MAP score of a library was the mean oriented contrast over MAPs with a measurable profile (8,027–8,033 per library), with the two orientation sets weighted equally, standardized by the mean and standard deviation of the 48 cancer specimens (Figure 4D). Snyder libraries, aligned to hg19, were scored at the hg19 positions.

Because 14 patients contributed more than one specimen, confidence intervals and P values came from resampling or permuting whole patients. Correlations with tumor fraction across the 48 cancer specimens and the 30 patient means used Spearman ρ, with patient-bootstrap confidence intervals and patient-label permutation (Figure S6A). For all 54 libraries (Figure 4D), tumor fractions were permuted between patients with the same number of libraries, all libraries of a patient moving together and each control forming its own block (two-sided; 10,000 permutations). Within-patient association was tested by centering values within each of the 14 patients with repeated specimens and permuting the centered tumor fractions within patient (10,000 permutations; Figure S6B). A partial Spearman correlation adjusted the patient-level association for estimated depth and median fragment size (Figure S6C,D), and single-MAP correlations were computed across the 30 patients (Figure S6E).

#### Deep plasma WGS: representative locus and shift of peak protection

The locus in Figure 4C was chosen among K4/TF-correlated and K9/TF-correlated MAPs as one at which specimens in the highest tumor-fraction quartile had more protection than both the lowest quartile and BH01 at the cancer-associated position and less at the other position; library profiles were normalized and smoothed as above before averaging.

In each library, the profiles of the 1,776 K4/TF-correlated and the 1,619 K9/TF-correlated MAPs (Figure 4E, F), were divided by their ±250-bp window mean, oriented with the H3K9me3-associated position on the negative side, averaged into two composites, smoothed as above and scaled to the peak for display (Figure 4E). For each library, ranked by tumor fraction, the K9/TF-correlated composite was subtracted from the K4/TF-correlated composite, and the peak shift was the K4/TF-correlated minus the K9/TF-correlated peak position, searched between −60 and +30 bp (Figure 4F).

The two groups were also redefined among all 35,120 MAPs at |ρ| > 0.1, 0.2 and 0.3 (4,435 and 3,822; 1,776 and 1,619; 681 and 695 MAPs). In each specimen, the K9/TF-correlated mean contrast was subtracted from the K4/TF-correlated mean, and the difference was correlated with tumor fraction across the 48 cancer specimens and the 30 patient means (Spearman ρ; 2,000 patient-label permutations; Figure S6F).

#### Deep plasma WGS: sequencing depth, number of MAPs and in silico dilution

Separation of specimens in the lowest (tumor fraction 17–45%) and highest (51–82%) quartiles was measured after reducing depth or the number of MAPs. Reads were downsampled from 75-fold to 30-, 10-, 5- and 2-fold by keeping each read pair with probability equal to the target divided by the estimated depth, and at full depth 20 to 500 MAPs per direction were used. In each repeat, one half of the specimens ranked MAPs by quartile AUC and fixed their direction, and the other half was scored; test scores from all repeats were pooled into one ROC curve per setting (Figure S7A,B). Control panels used 500 loci per direction drawn from discovery-paired loci whose two marks did not change in opposite directions with tumor fraction.

Figure S7C compared paired-position, H3K4me2-position-only and H3K9me3-position-only measurements at 10–1,000 selected loci over 100 specimen partitions (24 training and 24 test specimens), with loci and directions selected in training data. Figure S7D,E compared, at each downsampled depth, models built on all MAPs, on K4/TF-correlated plus K9/TF-correlated MAPs or on either group alone over 100 partitions of 24 specimens from 21 patients (twelve training and twelve test specimens), with preprocessing and tuning in training data.

Lower tumor fractions were simulated by in silico mixing (Figure S7F). The raw profile (before normalization and smoothing) of one of two cancer specimens (tumor fraction 82% and 77%) and that of one Snyder library, each divided by its number of 120–180-bp fragments, were weighted to target tumor fractions of 0–20%, and each mixture was normalized, smoothed and scored as one library. For these mixtures, the hg19 Snyder libraries were placed in the hg38 frame, reversing 16 loci on the opposite hg38 strand and excluding 2 with build-dependent spacing.

#### Hybrid-capture sequencing

Emory capture reads were aligned to hg38 with BWA-MEM (-M; v0.7.17),^72^ duplicates were removed with Picard MarkDuplicates (v2.27.5), and reads in the ENCODE hg38 exclusion list (v2)^75^ were removed. Fragments were reconstructed from read 1 of properly paired, same-chromosome primary alignments with non-zero template length and mapping quality ≥30, excluding duplicates and quality-control failures. The WPS at each position was computed from fragments of all lengths, divided by the number of fragments overlapping ±250 bp and expressed per 100 fragments.

For Figure 5A, the 8,044 MAPs of the deep WGS score and their orientations were carried over unchanged, and the score was computed over the 224 MAPs with both positions inside the capture targets, so that no Emory data were used to choose loci or directions. The specimen score was the mean oriented MAP contrast over these MAPs with at least ten fragments at both positions, the measurability rule of the other capture analyses (median 200 MAPs per specimen; range 177–202). Serial specimens were summarized by the median within patient, and patients with mCRPC were compared with controls by two-sided Mann–Whitney test and by AUC with paired patient-bootstrap confidence intervals (20,000 resamples, stratified by group). For Figure 5B, two loci, one at which protection is expected to shift toward the H3K4me2-associated position with tumor DNA and one toward the H3K9me3-associated position, were chosen as those with the largest coverage-weighted difference in the expected direction between patients with mCRPC and controls, and were compared between the specimens of the two groups by patient-label permutation. The capture score of Figure S8 used 176 MAP candidates within 500 bp of a capture target whose discovery H3K4me2 and H3K9me3 signals correlated with tumor fraction in opposite directions, with ρ > 0.2 for the rising mark (63 with H3K4me2 and 113 with H3K9me3 rising); no Emory data were used to choose them. The score was the mean H3K4me2-minus-H3K9me3 contrast over the first set minus that over the second, using loci with at least one fragment at both positions in each specimen.

#### Tumor-fraction models in Foundation Medicine data

One prostate specimen per patient (89 eligible specimens from 80 patients) was chosen by de-identified specimen identifier, as analysis dates were unavailable, and patients were split into training (48), development-validation (16) and test (16) groups. Of the MAPs covered by the panel, 1,342 had at least ten 120–180-bp fragments at both positions in at least 16 of 20 specimens chosen without tumor-fraction labels, at evenly spaced ranks of BAM file size among the 1,285 eligible specimens. WPS at each position was divided by the number of 120–180-bp fragments overlapping ±73 bp and expressed per 100 fragments. Each MAP predictor was the MAP contrast, set to missing when either position had fewer than ten fragments, and specimens with at least 80% of predictors available, assessed on the original feature set, were kept (46, 15 and 16 patients; Figure S9A). Copositioned anchors gave 412 predictors in the same way.

A comparator model used 16 prespecified features (Figure S9H) modeled on the fragment-length and coverage features of APEX:^9^ seven for fragment length (10th, 25th, 50th, 75th and 90th percentiles, mean counted length and the fraction of fragments of 120–180 bp),^2,3^ five for sequencing quality (log mapped and unmapped reads, and paired, properly paired and duplicate fractions) and four for coverage at the sites the MAP model reads (the fraction of loci with at least ten fragments at both positions among the 1,342 MAPs, 581 H3K27me3/H3K36me3 position pairs from the inter-laboratory replicates, 412 copositioned anchors and 50 matched Non-MAP loci). A further comparator used sequencing depth (log mapped reads) alone.

Elastic-net regressions^85^ of reported tumor-fraction percentage used median imputation and standardization estimated within training folds. Because the 16 test patients had been used in an earlier analysis, discrimination was estimated by nested cross-validation in the 59 development patients with reported tumor fraction 0 (30) or ≥3% (29): 46 training and 13 development-validation patients, the other two development-validation patients having tumor fraction between 0 and 3% (Figure 5C). The five outer folds, stratified by class, were identical for all models (MAPs, copositioned anchors, the 16 features, MAPs plus the 16 features, and depth), fitted as elastic nets (Figure S9I) or ridge regressions (Figure S11C–E), with hyperparameters tuned by stratified five-fold cross-validation within each outer training set. AUC differences came from paired stratified patient bootstrap (2,000 resamples). In the test set (Figure S9B,C), models fitted in the 46 training patients were applied without refitting to the 16 test patients; agreement with reported tumor fraction was summarized by Spearman ρ, and the primary comparison was the paired difference in absolute error between MAP and copositioned predictions bounded to 0–100%.

In breast (389 patients) and lung cancer (378 patients), one specimen per patient was chosen by specimen identifier and predictors were computed as for prostate. Of these, 360 and 341 patients had at least 80% of MAP predictors, assessed on the original feature set, and inputs for every model, and 336 and 322 had reported tumor fraction 0 or ≥3% (218 and 118; 260 and 62); the 24 and 19 patients between 0 and 3% were not used. Models were refit within each cancer by nested cross-validation with five outer and five inner folds, stratified by class and shared by all models, with imputation, scaling, hyperparameters, thresholds and linear calibration estimated from outer-training patients (Figure 5H,I; Figure S9D–G,I,J; Figure S11D,E; Figure S12A–C). Errors and Pearson correlations (two-sided t-test) used patients with reported tumor fraction ≥3% (breast, 118; lung, 62), with confidence intervals from 1,000 patient-bootstrap resamples. In a fixed split of the same patients into training, validation and test groups (breast, 199, 67 and 70 patients; lung, 193, 63 and 66), models fitted in training patients, with thresholds and calibration from validation patients, were applied unchanged to test patients (Figure S12D,E).

Virtanen–Kwan models trained on the MAPs measurable on both panels (19 and 18 for the capture batch and the additional capture batch), with imputation and scaling learned in that cohort, were applied unchanged to the 59 Foundation Medicine prostate patients and compared with nested cross-validation on the same MAPs (Figure S11A,B). AUCs weighted each patient equally. Models were implemented in Python 3.11 with scikit-learn (v1.9.0).^86^

#### Tumor-fraction models in the Virtanen–Kwan cohort

Within each panel generation, models were fitted in development patients, the model size was chosen in tuning patients, and the chosen model was applied once to held-out patients. A MAP was measurable for a generation when development libraries had at least ten 120–180-bp fragments at both positions. Only the 3rd and 4th generations had at least 50 measurable MAPs (62 and 99). The 4th generation, with the most measurable MAPs and the most held-out patients (163), formed the capture batch (Figure 5), and the 3rd generation (58 held-out patients) formed the additional capture batch (Figure S10).

Elastic nets of the MAP contrasts were fitted with the top 25 or 50 MAPs, ranked by the absolute standardized coefficients of the full development model, or with all measurable MAPs, using patient-weighted mean imputation and standardization learned in development data and predictions bounded to 0–100%. The model size with the lowest patient-weighted mean absolute error in tuning patients was chosen before any held-out library was scored, with ties going to the smaller model: 50 of 99 MAPs for the capture batch and 50 of 62 for the additional batch (models on all 99, Figure S11A). Comparators fitted on the same development data were an elastic net on copositioned anchors, a regression on sequencing depth, and the training-set mean and median.

Selected models were applied without refitting or recalibration to held-out libraries with at least 80% of inputs measurable, and comparators were evaluated on the same libraries. AUCs compared reference tumor fraction 0 with ≥3% (or with each band in Figure 5E), excluding libraries between 0 and 3%, a threshold close to the detection limit of copy-number-based estimators;^42^ errors, correlations and calibration slopes (prediction on reference) used libraries with reference tumor fraction ≥3%. The operating point of Figure 5E was the predicted tumor fraction that maximized balanced accuracy (equivalently, Youden’s index) in tuning patients (16.1%). Each patient contributed a total weight of one, and confidence intervals were percentile intervals from patient-bootstrap resamples (1,000; 2,000 for the sensitivity and specificity in Figure 5E and for Figure S11A). Five patients contributed different draws to both held-out sets.

#### Serial draws

Serial pairs were consecutive draws of a held-out patient, ordered by draw number. Agreement between predicted and reference changes was summarized by patient-weighted Pearson correlation and by the patient-weighted percentage of pairs with the same sign of change among pairs with a reference change of at least 5 percentage points (other minimum changes, Figure S10E). For patients with at least three draws, the changes from the first to the second and from the second to the third draw were correlated in the same way (Figure 6D). Sensitivity analyses excluded pairs with reference tumor fraction 0 at both draws or kept only the first pair per patient (Figure S13A), and treatment history in Supplementary Table S1 of Virtanen, Kwan et al.^55^ was compared between the two draws of each pair (Figure S13B). P values for the patient-weighted correlations of Figures 5F, 6A and 6D came from two-sided permutation tests (10,000 permutations) that exchanged reference values between patients with the same number of libraries or changes, all values of a patient moving together.

#### Comparison with Fragle

Fragle (https://github.com/skandlab/FRAGLE, commit 3212220)^43^ was run in targeted-sequencing mode with its pretrained low- and high-tumor-fraction networks and model selector, using the genome build of each BAM header (hg38 for the Virtanen–Kwan cohort and hg19 for Foundation Medicine). As panel bait files were not available, target masks were inferred from coverage in 20 development patients of each cohort, one library per patient (Foundation Medicine prostate training specimens): autosomal 1-kb bins were retained when their depth was at least 20-fold the background (the mean depth after removing the 1% of bins with the highest depth) in at least 16 of the 20 libraries, refined to 100-bp resolution with the same rule within the retained regions and 1-kb flanks, and merged without padding. Fragle read the fragments outside the mask. The Foundation Medicine mask was applied unchanged to breast and lung cancer, where an elastic net on the length distribution of the same off-target fragments (specimens with at least 1,000) was also fitted on the same patients and folds as the MAP models.

Classification thresholds maximized Youden’s index. Thresholds and linear calibration were estimated from the 235 tuning libraries (157 patients) of the Virtanen–Kwan cohort, with equal total weight per patient, or from outer-training (nested cross-validation) or validation patients (fixed split) in Foundation Medicine, and were applied unchanged. MAP models and Fragle were compared on identical libraries and serial pairs, with 95% confidence intervals for paired differences from 1,000 patient-bootstrap resamples (Figures 5G and 6C; Figure S12).

#### Lineage programs

ASCL1-site MAPs (667 loci) and copositioned anchors (175 loci) were loci with either position within 250 bp of an ASCL1 ChIP-seq peak in small-cell lung cancer cell lines (GSE69394),^87^ and recurrent-AR-site MAPs (96 loci) those with either position within 250 bp of an androgen receptor binding site present in at least 25 of 88 primary prostate tumors (GSE120738);^88^ the 13 MAPs and two copositioned anchors at both kinds of site were excluded. For each locus, the H3K4me2- and H3K9me3-associated WPS values were each divided by the fragment count at that position and multiplied by 100, and the second was subtracted from the first. The patient score was the mean over loci with a fragment at each position (at least ten loci), and the prespecified primary score was the ASCL1-site minus the recurrent-AR-site score.

In the Baca cohort,^20^ PRAD and NEPC were compared by AUC with patient-stratified bootstrap confidence intervals (20,000 draws) and one-sided permutation tests in the direction expected for each lineage (100,000 label permutations), with Holm adjustment across the ASCL1-site and recurrent-AR-site scores (Figure 6F–I). A set of 220 ASCL1-site MAPs at peaks present in all four prostate NEPC ASCL1 ChIP-seq profiles (GSE156290) was analyzed in the same way. In the DFCI NEPC-monitoring cohort, specimen scores were summarized by the median within patient and state, and in the six patients with both states the two states were compared by AUC and by one-sided exact permutation of state labels within patient (Figure 6J, left). The first earlier and the first NEPC-labeled draw of each of these patients were compared by exact one-sided sign test for the MAP score, the copositioned score and reported tumor fraction (Figure 6J, right).

### Statistics and reproducibility

Sample sizes were set by the specimens available; no sample-size calculation was performed. No intervention was assigned, and investigators were not blinded to clinical labels. Serial specimens were analyzed within patient or summarized to one value per patient, and resampling was done by patient wherever a patient contributed more than one specimen. Tests were two-sided except where the hypothesis set the direction: larger separation at MAPs than at copositioned anchors (Figure 2C), retention of the discovery order of the two marks (Figure 2E) and the expected direction of the lineage scores (Figure 6F–J). P < 0.05 was considered significant. Multiple comparisons were adjusted within prespecified families: the four MAP-group-by-cohort comparisons (Holm; Figure 2C), the 59 reference region sets (Benjamini–Hochberg; Figure 3B,C), the factors of the separation test (Holm; Supplementary Table 3) and the two single-site lineage scores (Holm; Figure 6F–I). Analyses were performed in R and Python with the software versions given in each section.

## DATA AVAILABILITY

Plasma cfChIP-seq, targeted cfDNA and matched clinical data of Sipola et al., including the Wyatt multi-mark cfChIP-seq libraries used as inter-laboratory replicates, are available under controlled access through dbGaP (phs003482.v2.p1). Capture-panel data of Virtanen, Kwan et al. are available under controlled access from the European Genome-phenome Archive (EGAD50000001594). Deep WGS data of Herberts et al., control WGS of Snyder et al. and low-pass WGS of the Baca cohort are available as described in those studies. Public reference data used here are available from EpiMap (BSS01456), SCREEN v4, ENCODE (ENCSR315NAC, ENCFF685MPH, ENCFF001TDO, ENCSR000EGM, ENCSR000AKB), GTEx and the Gene Expression Omnibus (GSE69394, GSE120738, GSE43720 and GSE156290). Hybrid-capture data of the Emory capture cohort are not publicly available; fragment counts and protection scores at the analyzed MAP positions are available from the corresponding author. Low-pass WGS of the DFCI NEPC-monitoring cohort is described in G. G. Lee et al. (manuscript in preparation) and is not released with this study. Foundation Medicine data are not released due to privacy considerations, but can be requested under a data transfer agreement. cfChIP-seq data of the discovery cohorts are available from the corresponding author upon reasonable request. MNase–ChIP-seq data of LNCaP cells are available from the corresponding author upon reasonable request.

## CODE AVAILABILITY

cfChIP-seq preprocessing used SNAPIE (https://github.com/prc992/SNAPIE; versions in Methods).^69^ Code for MAP construction, fixed-position WPS extraction, statistical analysis and figure generation will be released at publication (https://github.com/yukunou703/MAPs).

## SUPPLEMENTARY INFORMATION

**Supplementary Note 1. Distance rules for pairing positions within cohorts and matching them across cohorts**

Pairing within a cohort was limited to 73 bp, approximately half of the 147-bp nucleosome core particle.^48^ Two dyads d bp apart share 147 − d bp of wrapped DNA: at d = 73 bp they share 74 bp (50.3%), and at 74 bp they share 73 bp (49.7%). The 73-bp limit is therefore the largest separation at which the two positions still share more than half of the core, a threshold for comparing alternative, overlapping nucleosome footprints across profiles. At 74 bp, the inferred footprints still overlap, but by less than half of the core. With nucleosome repeat length L, a dyad is nearer the neighboring nucleosome than nucleosome n once d exceeds L/2. Because L cannot be below 147 bp, L/2 is at least 73.5 bp; for the 167-bp plasma mononucleosomal fragment mode it is 83.5 bp, and for typical repeat lengths of 185–200 bp it is 92.5–100 bp. The 73-bp limit therefore restricts the analysis to strongly overlapping nucleosome footprints (**Figure S1F**).

We then examined how the distance windows affected pairing within discovery cohorts and matching between cohorts (**Figure S1D,E**; Supplementary Note Table 1). Two pairing rules were compared: the first assigned each H3K4me2-associated position to its nearest H3K9me3-associated position within the window (nearest-partner assignments); the second retained a pair only when the two positions were each other’s nearest partners (reciprocal pairs). At a 73-bp window, the first rule produced only 0.46% and 1.23% more pairs than the second in the two cohorts; at 150 bp, the differences were 8.89% and 11.22% (Supplementary Note Table 1a). With the within-cohort window held at 73 bp, increasing the cross-cohort matching tolerance from 20 to 30 bp increased the number of matched loci from 91,471 to 120,489, and a 50-bp tolerance yielded 161,975 (Supplementary Note Table 1b). Among candidate pairs with at least 10-bp separation in both cohorts, agreement in the order of the two marks was 69.4%, 66.1% and 59.9% at 20, 30 and 50 bp. The 30-bp setting recovered more loci than 20 bp and kept more directional agreement than 50 bp. At this tolerance, 35,120 of 53,131 direction-eligible candidate pairs had the same order in both cohorts (66.1%) and 18,011 had opposite orders; MAP selection required concordant order.

## Supplementary Tables

**Supplementary Tables 1–8** will be provided with the peer-reviewed version and are available from the corresponding author on request.

**Supplementary Table 1.** Cohorts, assays, tumor-fraction estimators and thresholds used in each analysis.

**Supplementary Table 2.** Chromatin-state, gene-feature, repeat, binding-site and candidate cis-regulatory element comparisons of MAPs with copositioned K4/K9 anchors, including displacement strata, the depth-stratified comparison of 59 reference region sets (**Figure 3B–D**) and the same comparison with anchors defined at 2, 4 and 8 bp.

**Supplementary Table 3.** Separation-gradient tests: odds ratio per 10-bp separation, 95% confidence interval, *P* value and Holm-adjusted *P* value for every factor under the eight declared conditions, with the per-factor classification and per-bin binding percentages.

**Supplementary Table 4.** CTCF proximity by peak strength and LNCaP ABC candidate-element and enhancer–gene link comparisons.

**Supplementary Table 5.** Tumor-fraction-associated MAP groups (K4/TF-correlated, K9/TF-correlated and neutral): locus lists, host-gene and bound-factor comparisons.

**Supplementary Table 6.** Virtanen–Kwan and Foundation Medicine per-library predictions.

**Supplementary Table 7.** Serial pairs: library identifiers, reference and predicted tumor fraction and treatment-history fields used for the draw-order check.

**Supplementary Table 8.** Breast and lung cancer refit by nested cross-validation and in a fixed split, and shared-MAP transfer between prostate cohorts: per-patient predictions.

